# Generalized calibration across LC-setups for generic prediction of small molecule retention times

**DOI:** 10.1101/2020.01.14.905844

**Authors:** Robbin Bouwmeester, Lennart Martens, Sven Degroeve

## Abstract

**Motivation:** Accurate prediction of liquid chromatographic retention times from small molecule structures is useful for reducing experimental measurements and for improved identification in targeted and untargeted MS. However, different experimental setups (e.g. differences in columns, gradients, solvents, or stationary phase) have given rise to a multitude of prediction models that only predict accurate retention times for a specific experimental setup. In practice this typically results in the fitting of a new predictive model for each specific type of setup, which is not only inefficient but also requires substantial prior data to be accumulated on each such setup.

**Results:** Here we introduce the concept of generalized calibration, which is capable of the straightforward mapping of retention time models between different experimental setups. This concept builds on the database-controlled calibration approach implemented in PredRet, and fits calibration curves on predicted retention times instead of only on observed retention times. We show that this approach results in significantly higher accuracy of elution peak prediction than is achieved by setup-specific models.

## Introduction

Mass spectrometry (MS) coupled to liquid chromatography (LC) is a key method for high-throughput analysis of the metabolome. The LC-based separation, which separates analytes based on their broader physicochemical properties, is carried out before the MS analysis, and ensures that only a fraction of the analytes compete for ionization over time, leading to less isobaric analytes being captured in the same fragmentation spectrum. LC separation thus enables more sensitive identification of low abundant analytes, as there is less competition for ionization, and as isobaric analytes are more likely to result in individual fragmentation spectra^1–3^. In addition to these benefits, the retention time (*t*_*R*_) of an analyte provides complementary information to the mass-to-charge (m/z), as it derives from a broader set of physiochemical properties of the analyte. This complementary information can be especially beneficial in metabolomics where many of the analytes are isobaric ^4,5^.

Even though the retention time has been shown to be a useful component for the identification of analytes ^4,6–17^, the incorporation in identification software remains limited. This is mainly due to the limited availability of retention time information in small molecule libraries, which in turn is tied to the variance in retention time caused by specific LC setups^8,18,19^.

It would therefore be ideal to be able to predict observed retention times on a given LC setup for all known small molecule structures in databases, which has resulted in increasing interest in modeling chromatographic setups and associated retention times. There are two main strategies to achieve this: retention time inference using observed retention times for a given set of analytes on different experimental setups as anchors ^4,20–22^, or predicting retention times from structure alone^4,6,8,23^. Because this first strategy relies on data from multiple setups for the same set of analytes, it requires that these analytes have been consistently observed across setups, and is limited by the number of different setups for which these analytes have been observed^24^. The second strategy finds the relation between structural features (e.g. quantitative structure-retention relationships^25^) and retention time using Machine Learning (ML) algorithms. Because of their predictive nature, these models are not limited by prior observations of an analyte, but rather by the availability of structures for the analytes of interest. However, this limitation is strongly mitigated due to the availability of extensive databases of small molecule structures ^26–28^.

As a result, such ML models have already been applied in non-targeted mass spectrometry to improve identification rates. For example, predicted retention times were used to halve the number of candidate isobaric lipids while retaining the majority of correct identifications^4^. In addition to limiting the search space, retention time predictions have also been used to decrease the number of false identifications for small molecules (< 400 Da)^11^, natural products from *Streptomyces*^12^, and sphingolipids^29^. While these methods are typically implemented down-stream of the identification process, an approach for the direct incorporation of retention time predictions in the identification process proper has also been developed^8^.

Nevertheless, these structure-based prediction models usually remain tied to a specific experimental setup, and perform very poorly for most other setups. This because differences in LC setup will significantly influence the retention times of analytes in complex ways, which results in non-transferable prediction models between setups. Even though the elution order is often conserved when the same type of column is used, there can still be dramatic variations in the retention times due to other differences in LC setup (e.g. in the RIKEN and FEM_long data sets as used in PredRet^18^). In practice, these differences therefore typically result in the fitting of a new predictive model for each experimental setup, even when there are only seemingly small differences in the setup. This in turn gave rise to a multitude of prediction models that only predict accurate retention times for a specific setup^8,30^.

A possible solution is provided by calibration between experiments, but current approaches for such calibration are mostly limited by matching observed retention times of analytes between the originally modeled LC setup, and the new LC setup. Importantly, however, this also means that generalization is lost, which means that accurate predictions for the new setup are now limited to only those analytes that were observed in the original setup. An example of this approach is PredRet^18^, which calibrates retention times between different experimental setups using Generalized Additive Models (GAM)^31^. In addition to calibration, an ML approach to predict the elution order of analytes has been developed based on the conserved elution order for specific column types across different LC setups^8^. However, the prediction of rank does not provide the same level of granularity as the prediction of exact retention time, and also requires specialized methods to incorporate in downstream analyses such as identification.

It can thus be clear that, despite very promising efforts to overcome the problem of across-setup retention time prediction, the problem has not yet been fully solved. Indeed, ideally we would be able to utilize the vast amount of data available in public repositories like MetaboLights^32^ and MoNA (http://mona.fiehnlab.ucdavis.edu/) to predict the retention time of any desired analyte on any kind of LC setup based on that analyte’s structure alone. This is all the more interesting as the combination of data from across many different experiments should provide more accurate predictions, and better generalization across a wider range of small molecules^6^.

We here therefore combine the two approaches of calibration and of generalization through ML to obtain a much more generic method to predict analyte retention times across LC setups based on structure alone. The result is our CALLC (CALibrate ALL LC) method, which uses a generalized calibration approach based on the mapping of retention time predictions between different LC setups. Interestingly, our approach also increases the amount of available data that can be used to fit the model, which in turn increases the predictive power of the model^8,30,33^.

## Methods

### Overview of CALLC

The objective of CALLC is to compute a retention time (*t*_*R*_) prediction model for a given LC setup from a number of data sets that contain observed analytes’ retention times, many of which can come from different LC setups. The goal of CALLC is therefore to generalize and calibrate previously trained predictive models from different LC setups for a specific LC setup. CALLC achieves this using three connected processing layers that each have their own distinct function (Figure 1).

**Figure 1:**
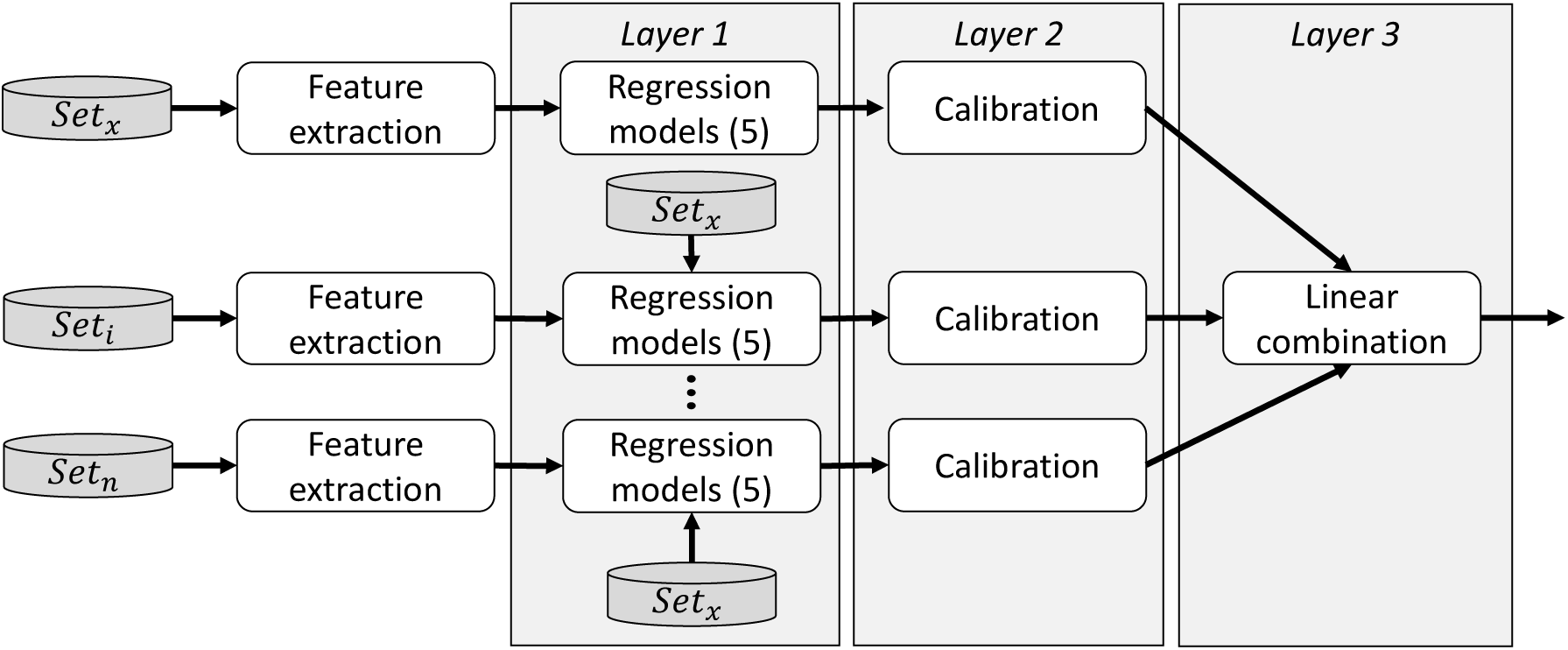
CALLC workflow using multiple models originating from different experimental setups. Each numbered data set derives from a given LC setup. For each data set, structural features for every molecule, and five setup-specific regression models are then trained in Layer 1. Predictions for the data set of interest (set_x_) from Layer 1 are then calibrated to the setup of interest (LC_x_) in Layer 2, and these calibrated predictions are then combined linearly in Layer 3 to yield a single predicted retention time per analyte.

The first layer implements the predictive model training approach in which a machine learning model is optimized for a specific LC setup (*LC*_*i*_), based on retention times obtained on that setup (*Set*_*i*_). CALLC uses five distinct regression algorithms to fit this model for the given LC setup. These five distinct algorithms are used because *a priori* selection of the best performing algorithm is decidedly non-trivial, and because combining multiple models actually improves prediction accuracy^30^. After training specific models (*M*_*i*_) for each specific LC setup (*LC*_*i*_), five *t*_*R*_ predictions *per* analyte are derived from each model *M*_*i*_ for the specific data set (*Set*_*x*_) obtained on the LC setup of interest (*LC*_*x*_).

The second layer calibrates all of these predictions for *Set*_*x*_ based on a similar approach to that of PredRet^18^. The key difference is that our approach uses predicted retention times instead of the observed retention times used in PredRet. The output of this second layer therefore again consists of five *t*_*R*_ predictions for each model *M*_*i*_ *per* analyte in *Set*_*x*_, but all these predictions have now been calibrated for setup *LC*_*x*_.

The third layer then linearly combines these calibrated *t*_*R*_ predictions from *Layer 2* into a single predicted retention time per analyte. Each layer is described in more detail below.

### LC setup specific data sets

A total of 42 experimental data sets containing a grand total of 4633 analytes from different experimental setups and labs were compiled from MoNA (http://mona.fiehnlab.ucdavis.edu/), PredRet^18^, and Aicheler^4^. After filtering duplicate analytes based on their InChI key a total of 2454 unique analytes remained across these 42 data sets. This compiled data set contains molecules with diverse molecular weights (from 44.078 Da to 2406.648 Da) and structures (from acetamides to lipids). Various data set properties, and their respective LC column types, are available in Tables S-1 – S-3.

### Features

RDKit is used to convert each InChI to numerical representations of the structure in what is called a feature vector^34^. A total of 196 features were calculated, which were filtered down to 157 features based on the requirement that each feature should have a standard deviation across analytes higher than 0.01, and squared Pearson correlation between features lower than 0.96. The original 196 features are listed in Table S-4, and the 157 filtered features are given in Table S-5.

### Layer 1

The first layer was trained using five machine learning algorithms: XGBoost^35^ (GB), Support Vector Regression^36^ (SVR), Least Absolute Shrinkage and Selection Operator^37^ (LASSO), Adaptive Boosting^38^ (AB), and Bayesian Ridge Regression^39^ (BRR). Every data set from Table S-1 was used to create its own set of five models. This yielded a total of 210 models for the 42 data sets. A ten-fold Cross-Validation (CV) with 25 randomly sampled hyperparameter sets was used for model optimization (see Code Listing S-1), because randomly selecting hyperparameters has been shown to require fewer iterations for optimization^40^. The hyperparameter set with the lowest Mean Absolute Error was used for training the model.

To calibrate predictions from an original LC setup to a new LC setup, CALLC needs training molecules with known *t*_*R*_ for the new setup (just as with PredRet^18^). These training molecules will be referred to as the calibration analytes. First, these calibration analytes are used to train five specific models for the LC setup of interest, and these models are added to the pool of pre-trained models from different LC setups. Second, predictions are made for the calibration analytes using all models in the pool. These predictions are made with the same cross-validation scheme that was defined for the hyperparameter optimization, which means that these calibration predictions are independent from the learned model parameters or hyperparameters. These predictions are then used as input for the second layer.

### Layer 2

The second layer takes the various predictions for the calibration analytes from *Layer 1* to fit a calibration curve that maps between the retention time predictions and observations. The calibration curve for the five newly trained models that were originally based on the calibration analytes constitutes the trivial case, and is therefore expected to be linear and have a slope of 1 and intercept of 0. In contrast, calibration curves for the other pre-trained models are expected to have a wide range of shapes: linear, sigmoidal, or even more complex functions.

These calibration curves are fitted using a Generalized Additive Models (GAMs) that uses thin plate splines from the R-package mgcv^41^. The GAM is able to fit a wide range of functions due to its additive nature, and is fitted for every model from *Layer 1* individually. The dimensions of the smoothing term are set to one degree of freedom (*k* − 1), while all other hyperparameters are kept at default values.

The cross-validation scheme from *Layer 1* is re-used to obtain predictions for the calibration analytes to avoid information leakage between the folds of the CV. The resulting calibrated predictions for the calibration analytes are subsequently blended in *Layer 3*.

### Layer 3

The calibrated predictions from *Layer 2* are here blended in a single prediction per calibration analyte using an elastic net^42^. This elastic net model is used to get a regularized linear combination of calibrated predictions that originate from different experimental setups and algorithms for the same analyte.

### Model evaluation

The CALLC architecture is evaluated using two analyses; layer performance and existing model comparison. The layer performance evaluation is repeated twice, once including duplicate analyte structures between data sets, and once with duplicate structures removed. This second evaluation tests whether differences in performance are solely due to the presence of duplicate structures. Interestingly, PredRet or similar calibration approaches would not be able to create a model without duplicate analytes across data sets.

The layers are evaluated with learning curves and a ten-fold CV strategy. The CV strategy is performed on two levels. On the first level, one fold is separated from the fitting procedure, with training in all layers based on the nine remaining folds. The separated fold, which is independent from parameter or hyperparameter optimization in any of the layers, is then used for final evaluation purposes across the three layers. Data sets are excluded from evaluation if they contain less than twenty analytes.

The learning curves use an increasing number of calibration analytes for training, with this number ranging from 20 to 100 in steps of 20 calibration analytes. The calibration analytes for each step are randomly sampled, and the remaining analytes are used for evaluation purposes. The whole procedure for each data set is repeated five times with different sets of calibration analytes. A data set is excluded from steps in the learning curve if less than ten analytes remain after selecting the calibration analytes. Importantly, each evaluation for the learning curve is based on a subset of the data that was not used in parameter or hyperparameter optimization in any of the layers.

The predicted retention times are evaluated between predicted and observed *t*_*R*_ using the Relative Mean Absolute Error (RMAE) and Pearson correlation (R) metrics. The MAE is made relative for each data set by dividing by the *t*_*R*_ of the last observed analyte. For evaluation of the layers, models from *Layer 1* and *Layer 2* are chosen to represent the layers based on the highest R or lowest RMAE on the training set CV. The exact metric is selected by matching it with the visualization metric for the test set.

The external comparison is made between *Layer 3* of CALLC and the reported performance from the Aicheler et al. SVR model^4^. Overlapping structures across data sets are allowed for CALLC.

## Data availability and study reproducibility

Scikit-learn^43^ V0.20.0, XGBoost^35^ V0.9, RDKit^34^ V2019.09.1 and Pandas^44^ V0.25.3 libraries for Python V3.6 were used. The library mgcv^41^ V1.8-31 was used for R V3.5.1. The code used to generate the regression models and make predictions and figures is available at:

https://github.com/RobbinBouwmeester/CALLC_evaluation

In addition to the code required to reproduce the research presented, CALLC has a user-friendly Graphical User Interface (Figure S-1) that is available at:

https://github.com/RobbinBouwmeester/CALLC

## Results

### Layer 1

The CALLC architecture consists of three connected layers (Figure 1). The first layer (*Layer 1)* uses a similar approach to fitting a conventional setup-specific *t*_*R*_ prediction model. The performance of each *Layer 1* algorithm for different numbers of calibration analytes is shown in Figure 2. In this comparison a representative model of *Layer 1* is selected by choosing the learning algorithm with the best CV performance (labeled as *Layer 1* in the figure).

**Figure 2:**
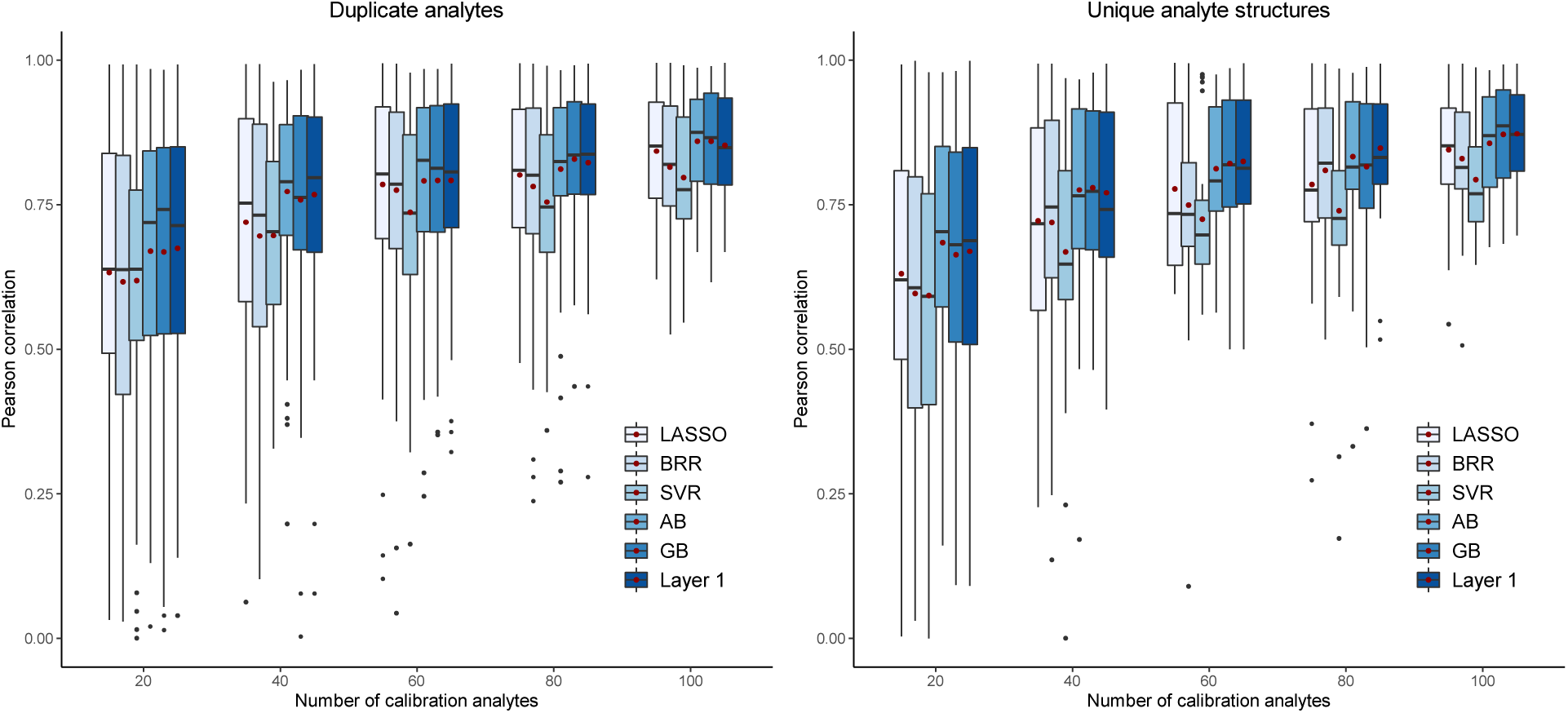
Comparison of different regression models in Layer 1 and the model predictions that were selected based on the CV performance (labeled as Layer 1). The evaluation metric is the Pearson correlation and the red dot shows the mean value of this metric. The left panel consists of 34 data sets that have shared analyte structures between data sets. The right panel consists of 21 data sets that do not share any analyte structures between data sets.

This comparison clearly shows that using more training analytes leads to higher performance, and as we have observed before^30^, there is not one algorithm that always outperforms the others. Instead of selecting a single learning algorithm, the best performing model is therefore used in further comparisons of the layers in CALLC.

There is no difference in performance between the two different analyses (with duplicate analytes between data sets and without duplicate analytes). For completeness, the same evaluation is repeated with the RMAE as the metric (Figure S-2), yielding identical conclusions.

### Layer 2

In the second layer (*Layer 2*) a GAM is used to calibrate the predictions from *Layer 1* for the experimental setup that is being evaluated. A GAM was chosen based on its relative simplicity and robustness to overfitting. Furthermore, a GAM is a suitable algorithm because a significant proportion of experimental setups have a conserved elution order, meaning that the calibration curve must be able to fit monotonal increasing or decreasing calibration curves. The capability of a GAM to fit complex calibration curves is shown in Figure 3 where the gradient profile of the solvents is different.

**Figure 3:**
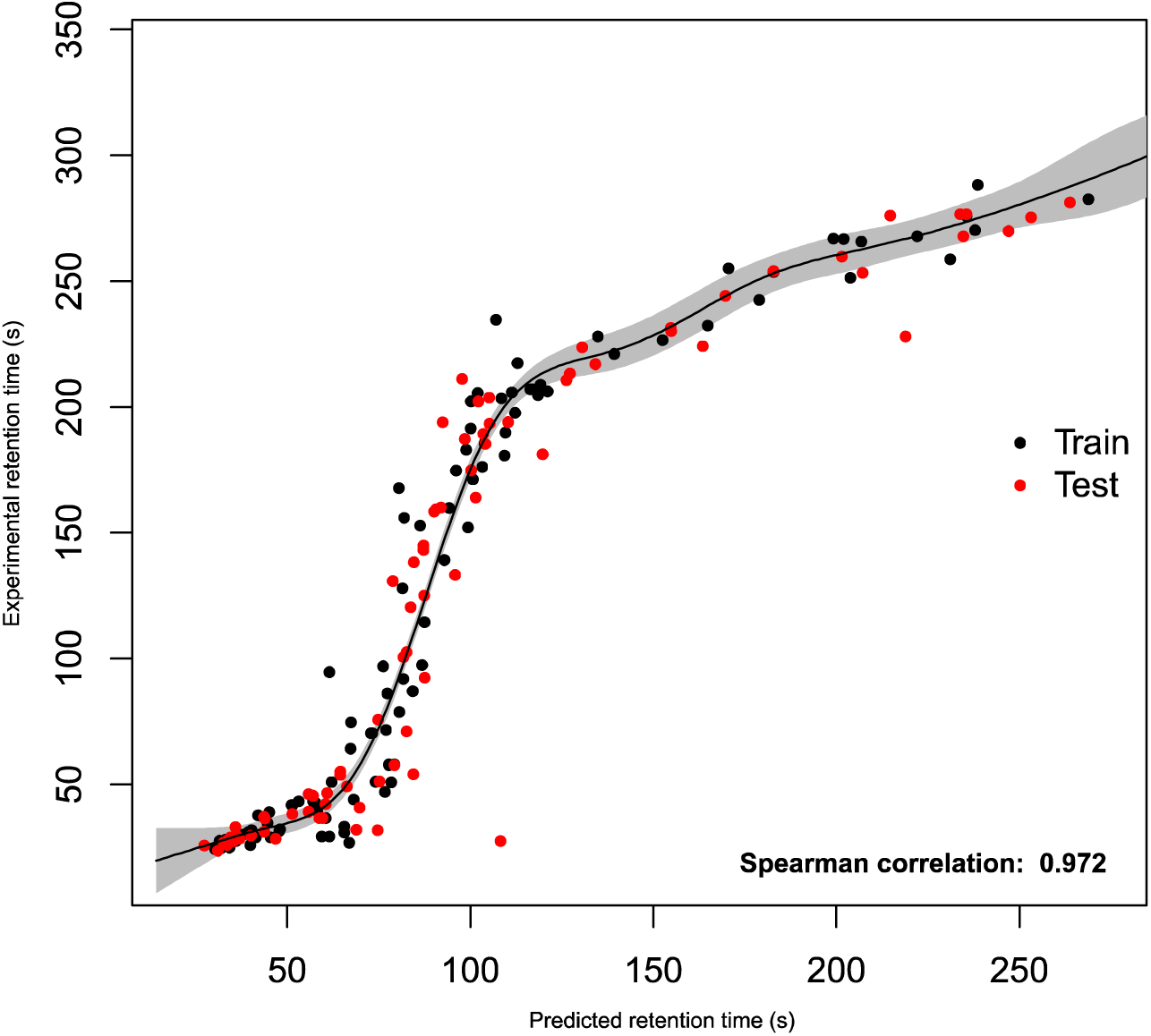
Example of a GAM model that is used to calibrate predictions from a model based on the ‘LIFE_old’ data set to the ‘LIFE_new’ data set. Black points show predictions used for fitting the calibration curve, while red points are part of the test set. The shaded grey area is the standard deviation of the fit.

The performance of the mapping mainly depends on the accuracy of predictions from *Layer 1*. Figure S-3 shows a less successful mapping due to inaccurate predictions from *Layer 1*. These kinds of inaccurate predictions, which remain inaccurate after calibration, do not contribute to an improvement of the prediction accuracy. In the third and last layer the calibrated predictions from *Layer 2* are combined into a single prediction *per* analyte, but in such a way that inaccurate (calibrated) predictions are likely to be ignored in this combination.

### Layer 3

After calibration in *Layer 2*, the final layer (*Layer 3*) of the model blends the predictions from different data sets and algorithms for a more accurate prediction. The learning algorithm in *Layer 3* should be able to handle the sparsity in (accurate) predictions from *Layer 2* to make accurate predictions, because a large proportion of calibrated predictions are not useful for achieving a higher performance (e.g. Figure S-3). An elastic net can do this because it blends predictions without overtraining due to regularization, and because of its relative simplicity of the trained linear model. An additional advantage of the elastic net is that the fitted coefficients, and the contribution of each model from the previous layers can be interpreted with relative ease.

### Layer evaluation

In this section, the performance of each layer is evaluated. The performance of *Layer 1* is based on the best performing model from *Layer 1*. Performance of this layer is determined by the model and data for the specific data set. The performance of *Layer 2* is based on the best CV performing model from *Layer 1* after calibration. Performance of this layer is determined by a single calibrated model selected from several data sets. Finally, for *Layer 3* no selection of models needs to be made, because a linear combination of all calibrated predictions is used. This also means that performance from *Layer 3* is not determined by single data sets or learning algorithms.

A significant difference in performance can be observed for the different layers in the learning curves (Figure 4). For both analyses, with duplicate analytes between data sets and without duplicate analytes, *Layer 3* achieves the highest performance for all the different number of calibration analytes. This is particularly noticeable for low numbers of calibration analytes (below 60). In addition, *Layer 2* outperforms *Layer 1*, especially when duplicate analytes are allowed across the data sets. This is unsurprising, because those analytes have been observed before and are therefore relatively easy to predict after calibration as shown before by PredRet^18^. However, when no overlapping analytes between data sets are available, calibration of predictions can still improve predictions. And even when there is no overlap in analyte structures the performance is increased by further combining these calibrated predictions in *Layer 3*. For completeness, the same evaluation is performed with the RMEA as a metric (Figure S-4), yielding the same conclusions.

**Figure 4:**
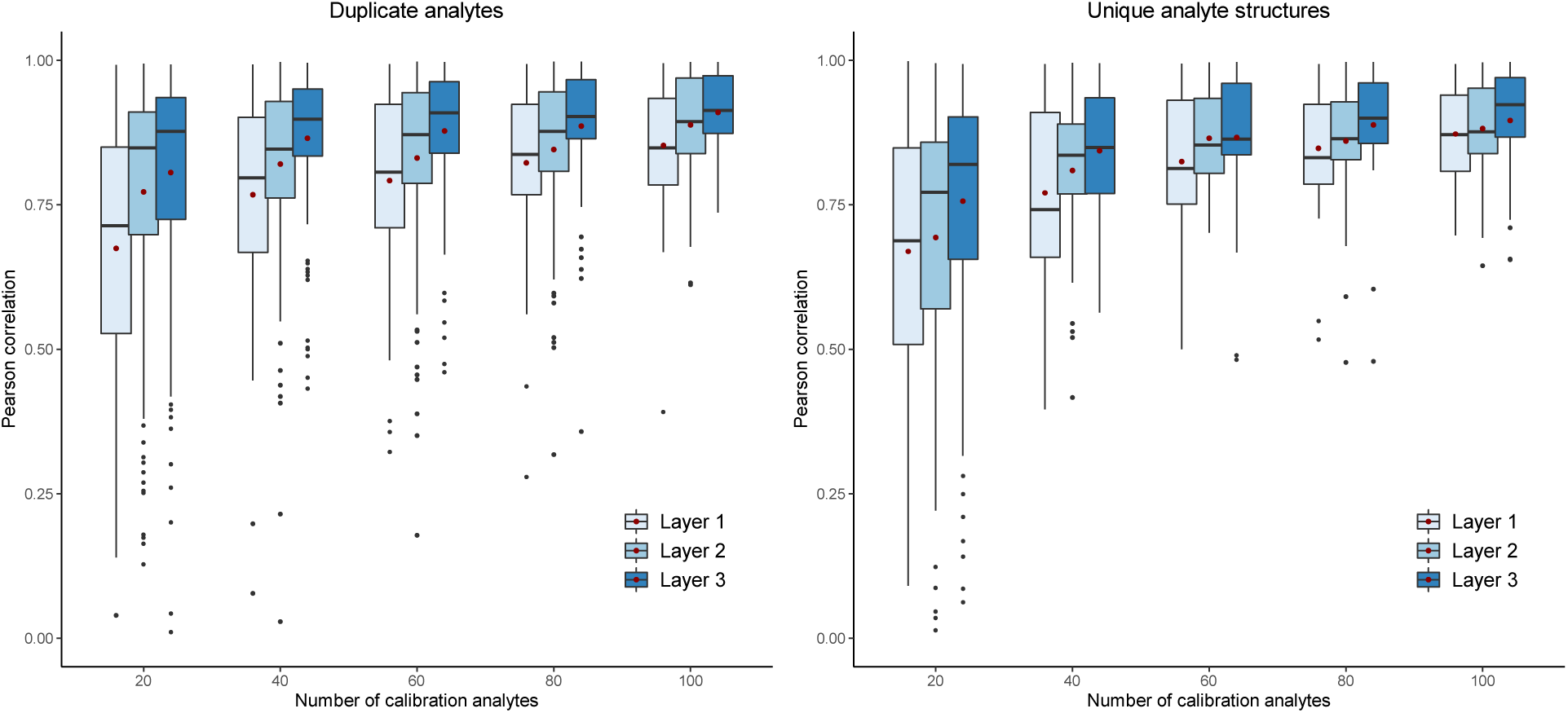
Performance comparison between the different layers using the Pearson correlation between predicted and experimental t_R_. In the left panel duplicated molecules are allowed for 34 data sets, while in the right panel duplicate molecules are removed for 21 data sets. The red dot shows the mean value.

An analysis with a ten-fold CV was used to show individual performance on the data sets (*Figure 5*). In this analysis, 16 out of the 40 data sets achieve an easily observable higher performance in *Layer 3* predictions compared to *Layer 1* predictions. Only a slightly better performance can be observed for 17 out of 40 data sets, and a slightly worse performance was observed for seven data sets.

**Figure 5:**
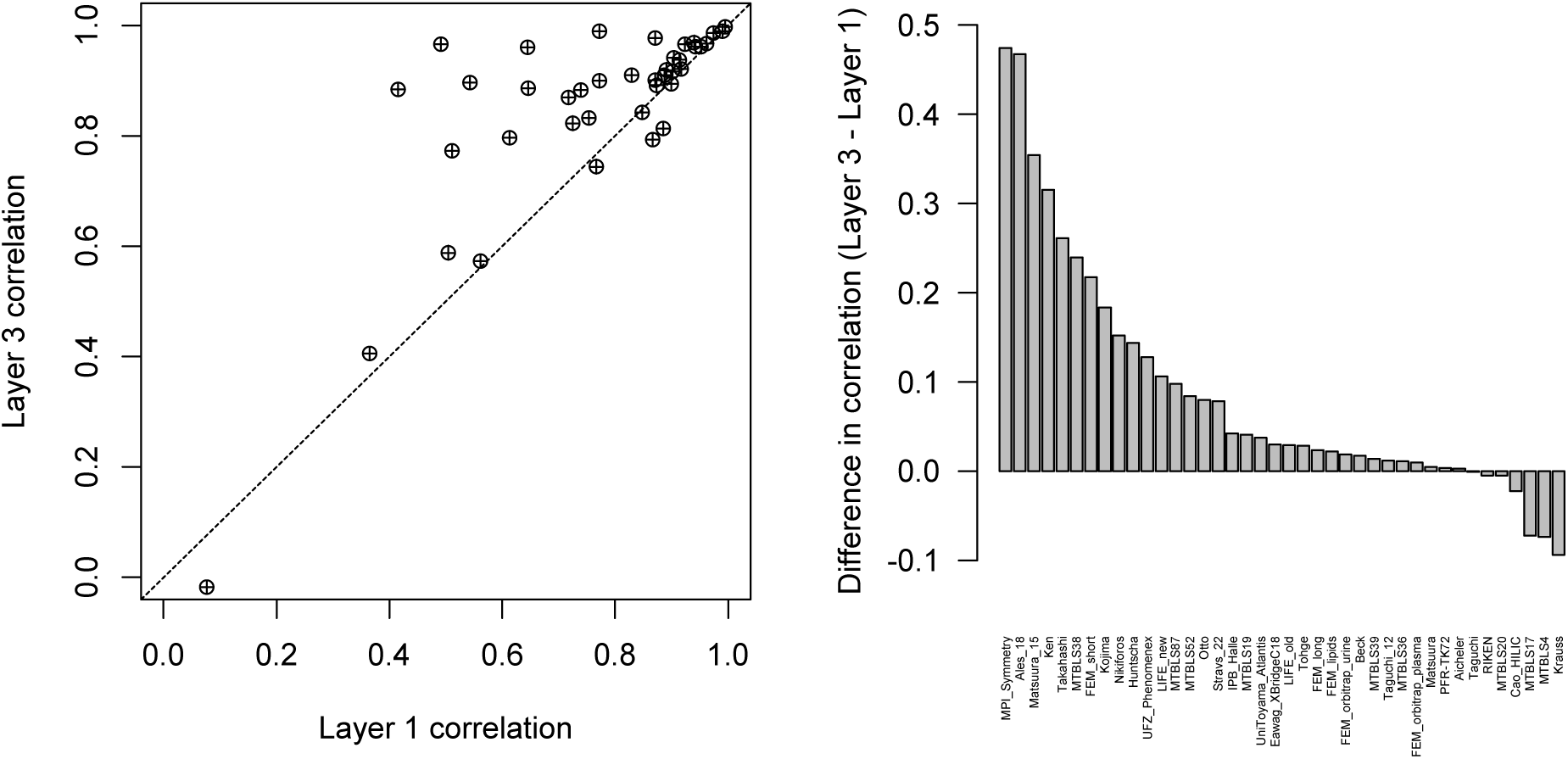
CV performance evaluation between Layer 1 and Layer 3 on 40 data sets, where shared analytes structures between data sets are allowed. The evaluation metric is the Pearson correlation between predicted and observed retention times. The left panel shows the achieved correlation for each data set in both layers, where the dotted line indicates the position where both layers perform equally. The right panel shows the difference in the Pearson correlation between the layers. Positive values mean that Layer 3 had a higher correlation than Layer 1, with the height of the bar showing the magnitude of the difference between the correlation values. Negative values show a higher correlation in Layer 1 than Layer 3.

Comparisons between *Layer 1* and *Layer 2* show that performance differences here are smaller (Figure S*-*5). Ten data sets achieve an easily observable higher performance for *Layer 2*, while the remaining data sets perform on par with *Layer 1*. For the comparison between *Layer 2* and *Layer 3*, almost all data sets had better or equal performance, except for two data sets that performed worse (Figure S*-*6).

For the analysis without duplicate structures between data sets the same observations are made (Figures S*-*7 – S*-*9). However, as expected, the difference in performance between *Layer 1* and *Layer 2* is smaller here due to the absence of overlapping analytes.

When the results from *Figure 5* are analyzed in more detail, it becomes clear why certain data sets show no improvement in *Layer 3* over *Layer 1* (Figures S*-*10 – S*-*13). Specifically, for the *Krauss* set, none of the layers show any real ability to predict analyte retention times. For *MTBLS4* and *MTBLS17*, their small data set sizes (less than 40 analytes) can potentially explain the worse performance. *Cao_HILIC* is only providing worse predictions for analytes that are non-retained (or at least that elude very early), because prediction performance of *Layer 3* is higher for longer retained analytes.

We can thus show that using several learning algorithms and incorporating more data increases the accuracy of retention time prediction for CALLC. Moreover, every layer in CALLC has its own distinct function, and all are critical to obtaining the highest possible performance. Importantly, these results show that overlap in analyte structures is not required to improve performance, and that the concept of generalized calibrations works well even when there is no overlap in structures between the data sets.

### Layer 3 coefficient interpretation

One of the advantages of using an elastic net in *Layer 3* is the relative ease of interpretation. These coefficients can be used to determine which prediction sets, from a specific learning algorithm and data set, are the most predictive for the data set of interest. These elastic net coefficients show only a slight clustering between used models (Figures S-14 and S-15), and, importantly, that *Layer 3* used a large variety of models to generate predictions for a data set.

### Comparison with the Aicheler model

To obtain an external evaluation of CALLC, a comparison is made with the SVR-based predictor from Aicheler et al.^4^ (Figure 6). This shows the added value of CALLC compared to existing strategies. In this comparison, CALLC demonstrates an average improvement of about 1.5 times for the MAE. Even when the procedure is repeated twenty times for each step, each time using different calibration analytes there are only two of the 340 rounds that perform worse than the MAE reported by Aicheler et al. While the difference between the models becomes smaller for large numbers of calibration analytes, CALLC performance remains significantly better.

**Figure 6:**
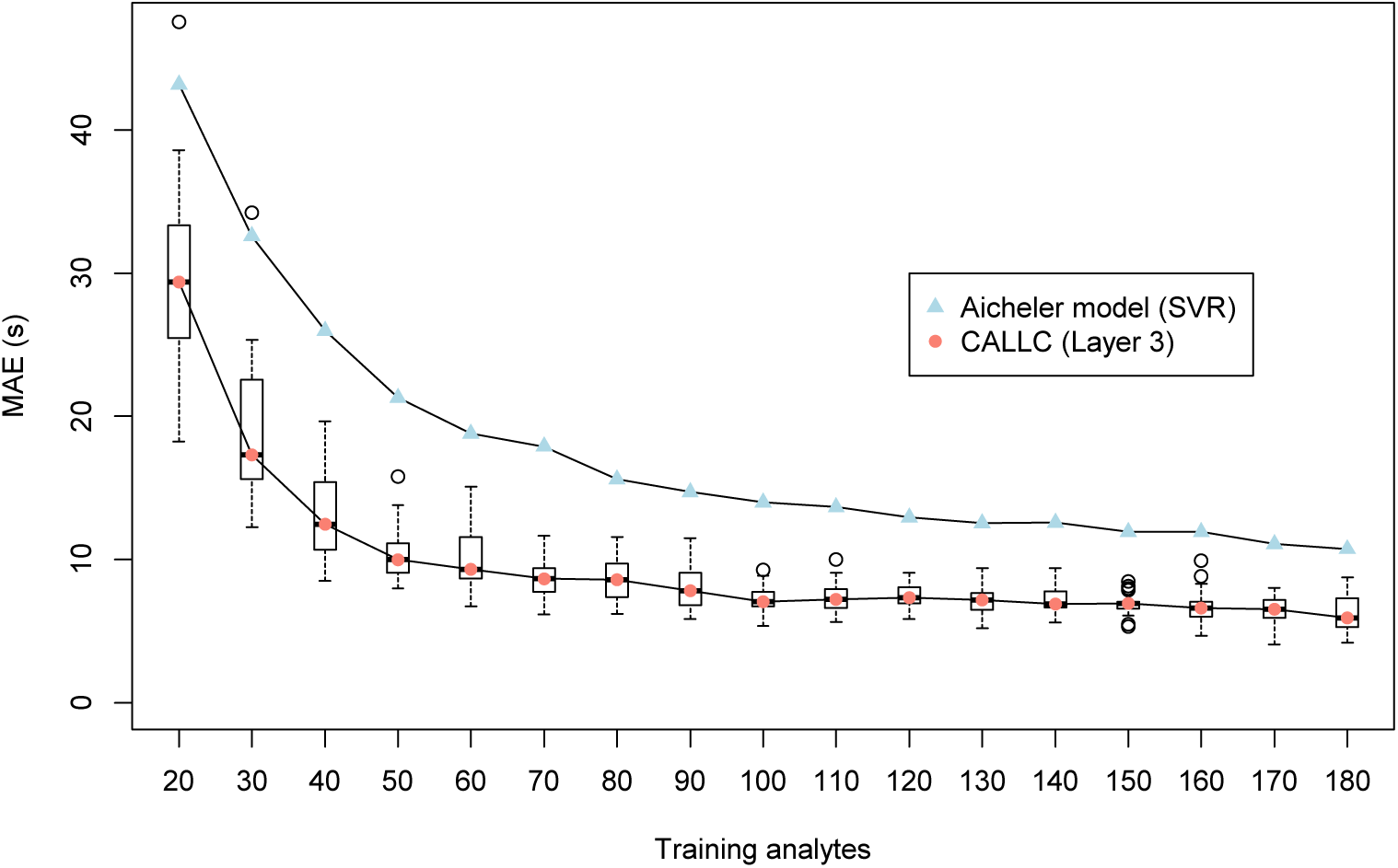
Performance comparison between an external t_R_ prediction model and this model. For CALLC the data sets contained duplicate structures across data sets. Error bars for different numbers of initial training instances are only shown for CALLC, while only average performance for the Aicheler et al. model could be obtained.

## Discussion

Retention time prediction has still not been used to its fullest potential in LC-MS, mainly because it is difficult to port predictions to different LC setups. To boost the use of retention time prediction, we here therefore introduced CALLC, which uses the concept of generalized calibrations for a more flexible application of retention time prediction and accurate predictions across LC setups. CALLC selects the most predictive molecular features, the most appropriate machine learning algorithms, and combines all information from individual pre-trained models. We show that using multiple data sets instead of a single data set improves prediction accuracy. Internal validation showed a significant increase in the performance of our approach, regardless of whether duplicated molecules were included (Figure 4 and 5). Moreover, external validation also shows a significant improvement in *t*_*R*_ prediction accuracy (Figure 6).

Of note, our strategy is adaptive because of its layered design. When a new data set is added, the model does not need to be retrained entirely. Indeed, a new model is only trained in *Layer 1* for the added data set. *Layer 2* and *Layer 3* are then very fast to retrain due to the single feature used in the calibration, and the inherent simplicity of the elastic net, respectively. The chosen learning algorithms or calibration method can also be swapped out to make the overall approach more suitable for any specific problems the researcher might be facing.

CALLC is also made freely available online as a software tool, which includes a Graphical User Interface to allow researchers to apply CALLC on their own data.

## Supporting information

All supporting Figures and Tables

