## Supplementary material for "Generalized calibration across LC-setups for generic prediction of small molecule retention times": All supporting Figures and Tables

Supplementary information for the paper entitled Generalized calibration across LC-setups for generic prediction of small molecule retention times.

Figure S-5: CV performance evaluation between Layer 1 and Layer 2 on 40 data sets, where shared analytes structures between data sets are allowed. The evaluation metric is the Pearson correlation between predicted and observed retention times. The left panel shows the achieved correlation for each data set in both layers, where the dotted line indicates the position where both layers perform equally. The right panel shows the difference in the Pearson correlation between the layers. Positive values mean that Layer 1 had a higher correlation than Layer 2, with the height of the bar showing the magnitude of the difference between the correlation values. Negative values show a higher correlation in Layer 1 than Layer 2. One data point, for Krauss, in the left panel is outside of the plotting area. ........................................................................................ 16

Figure S-6: CV performance evaluation between Layer 2 and Layer 3 on 40 data sets, where shared analytes structures between data sets are allowed. The evaluation metric is the Pearson correlation between predicted and observed retention times. The left panel shows the achieved correlation for each data set in both layers, where the dotted line indicates the position where both layers perform equally. The right panel shows the difference in the Pearson correlation between the layers. Positive values mean that Layer 3 had a higher correlation than Layer 2, with the height of the bar showing the magnitude of the difference between the correlation values. Negative values show a higher correlation in Layer 2 than Layer 3. One data point, for Krauss, in the left panel is outside of the plotting area. ........................................................................................ 17

Figure S-7: CV performance evaluation between Layer 1 and Layer 3 on 28 data sets, where shared analytes structures between data sets are removed. The evaluation metric is the Pearson correlation between predicted and observed retention times. The left panel shows the achieved correlation for each data set in both layers, where the dotted line indicates the position where both layers perform equally. The right panel shows the difference in the Pearson correlation between the layers. Positive values mean that Layer 3 had a higher correlation than Layer 1, with the height of the bar showing the magnitude of the difference between the correlation values. Negative values show a higher correlation in Layer 1 than Layer 3. One data point, for Krauss, in the left panel is outside of the plotting area. ........................................................................................................ 18

Figure S-8: CV performance evaluation between Layer 1 and Layer 2 on 28 data sets, where shared analytes structures between data sets are removed. The evaluation metric is the Pearson correlation between predicted and observed retention times. The left panel shows the achieved correlation for each data set in both layers, where the dotted line indicates the position where both layers perform equally. The right panel shows the difference in the Pearson correlation between the layers. Positive values mean that Layer 2 had a higher correlation than Layer 1, with the height of the bar showing the magnitude of the difference between the correlation values. Negative values show a higher correlation in Layer 1 than Layer 2........................................................................... 19

Figure S-9: CV performance evaluation between Layer 2 and Layer 3 on 28 data sets, where shared analytes structures between data sets are removed. The evaluation metric is the Pearson correlation between predicted and observed retention times. The left panel shows the achieved correlation for each data set in both layers, where the dotted line indicates the position where both layers perform equally. The right panel shows the difference in the Pearson correlation between the layers. Positive values mean that Layer 3 had a higher correlation than Layer 2, with the height of the bar showing the magnitude of the difference between the correlation values. Negative values show a higher correlation in Layer 2 than Layer 3. One data point, for Krauss, in the left panel is outside of the plotting area. ........................................................................................................ 20

*Table S-1: Properties of all 42 data sets used in the CALLC evaluation where duplicate structure across data sets are allowed.*

| Dataset | Identified molecules | Lowest MolLogP | Highest MolLogP | Lowest MW | Highest MW | Excluded in CV | Excluded in Learning Curve |
|---|---|---|---|---|---|---|---|
| Ken | 25 | -1.5149 | 4.6299 | 222.196 | 957.117 | | X |
| MTBLS17 | 24 | 1.851 | 5.5762 | 161.126 | 567.704 | | X |
| FEM_lipids | 70 | -1.8065 | 18.7691 | 144.214 | 891.501 | | |
| Taguchi | 38 | 12.0795 | 18.3211 | 663.037 | 907.459 | | |
| Takahashi | 75 | -3.8768 | 1.1223 | 103.121 | 809.578 | | |
| LIFE_old | 183 | -5.3972 | 11.4926 | 44.078 | 826.109 | | |
| UFZ_Phenomenex | 196 | -1.03416 | 6.718 | 113.12 | 837.058 | | |
| MTBLS52 | 24 | -3.8778 | 3.07084 | 386.353 | 860.771 | | X |
| Tohge | 77 | -2.892 | 3.3931 | 224.212 | 740.664 | | |
| MTBLS39 | 27 | -5.3956 | 1.5461 | 134.087 | 638.578 | | X |
| MTBLS87 | 146 | -5.3972 | 4.4779 | 105.093 | 911.667 | | |
| kohlbacher | 204 | 2.3565 | 13.6503 | 172.268 | 915.196 | | |
| MTBLS19 | 28 | -1.23283 | 8.9599 | 131.135 | 584.673 | | X |
| Krauss | 60 | -0.1953 | 6.2066 | 69.067 | 288.782 | | |
| Vogler | 5 | -1.03776 | 4.7235 | 102.097 | 314.469 | X | X |
| Matsuura | 83 | -20.484 | 1.9792 | 364.351 | 1828.691 | | |
| Taguchi_12 | 416 | 1.6921 | 39.6054 | 437.558 | 2406.648 | | |
| MTBLS36 | 84 | -7.5698 | 4.0662 | 88.062 | 504.438 | | |
| Huntscha | 22 | -1.2349 | 5.458 | 69.067 | 503.515 | | X |
| Ales_18 | 41 | -2.86343 | 5.134 | 137.182 | 822.953 | | |
| FEM_orbitrap_urine | 121 | -3.5854 | 7.8929 | 60.052 | 515.713 | | |
| Cao_HILIC | 108 | -5.6689 | 3.8257 | 59.068 | 663.43 | | |
| FEM_short | 72 | -3.0658 | 10.4322 | 74.079 | 906.94 | | |
| 1290SQ | 8 | -0.6459 | 4.3806 | 133.194 | 362.466 | X | X |
| LIFE_new | 173 | -5.3972 | 12.3886 | 75.111 | 834.173 | | |
| IPB_Halle | 76 | -2.44403 | 5.1358 | 112.088 | 521.676 | | |
| MTBLS20 | 136 | -5.1426 | 5.1358 | 60.056 | 521.676 | | |
| Otto | 39 | -1.98 | 6.1213 | 136.114 | 563.672 | | |
| Kojima | 35 | -3.3875 | 3.8441 | 175.187 | 513.504 | | |
| MTBLS4 | 34 | 3.7994 | 12.6126 | 467.584 | 836.189 | | |
| MPI_Symmetry | 41 | -2.86343 | 5.134 | 137.182 | 822.953 | | |
| Nikiforos | 68 | -1.0397 | 6.80734 | 114.173 | 916.112 | | |
| Beck | 270 | -3.0115 | 7.8493 | 74.083 | 1045.205 | | |
| MTBLS38 | 56 | -5.3972 | 5.3033 | 111.104 | 351.363 | | |
| UniToyama_Atlantis | 101 | -2.892 | 7.2336 | 222.196 | 1109.307 | | |
| Matsuura_15 | 30 | -20.484 | -6.753 | 545.491 | 1518.337 | | |
| Eawag_XBridgeC18 | 364 | -1.6561 | 11.919 | 119.127 | 995.189 | | |
| Stravs_22 | 119 | -1.0397 | 7.1837 | 129.159 | 785.891 | | |
| FEM_orbitrap_plasma | 120 | -2.4396 | 11.7242 | 77.152 | 786.129 | | |
| FEM_long | 443 | -9.7472 | 11.168 | 59.072 | 1272.453 | | |
| PFR-TK72 | 41 | -2.6973 | 7.2336 | 154.121 | 1034.2 | | |
| RIKEN | 351 | -7.573 | 7.5065 | 73.095 | 932.834 | | |

*Table S-2: Properties of all 42 data sets used in the CALLC evaluation where duplicate structure across data sets are removed.*

| Dataset | Identified molecules | Lowest MolLogP | Highest MolLogP | Lowest MW | Highest MW | Excluded in CV | Excluded in Learning Curve |
|---|---|---|---|---|---|---|---|
| Ken | 21 | -1.5149 | 4.6299 | 294.391 | 957.117 | | X |
| MTBLS17 | 16 | 1.851 | 5.4607 | 161.126 | 529.696 | X | X |
| FEM_lipids | 21 | 0.7317 | 18.7691 | 226.36 | 891.501 | | X |
| Taguchi | 37 | 12.0795 | 18.3211 | 663.037 | 907.459 | | |
| Takahashi | 53 | -3.8768 | 0.4444 | 103.121 | 809.578 | | |
| LIFE_old | 44 | -0.8663 | 11.4926 | 44.078 | 826.109 | | |
| UFZ_Phenomenex | 62 | 0.187 | 6.718 | 121.183 | 531.44 | | |
| MTBLS52 | 23 | -3.8778 | 3.07084 | 386.353 | 860.771 | | X |
| Tohge | 64 | -2.892 | 3.3931 | 224.212 | 740.664 | | |
| MTBLS39 | 7 | -2.2399 | 1.1307 | 332.261 | 624.551 | X | X |
| MTBLS87 | 49 | -5.2802 | 1.4621 | 145.202 | 911.667 | | |
| kohlbacher | 204 | 2.3565 | 13.6503 | 172.268 | 915.196 | | |
| MTBLS19 | 14 | -1.23283 | 8.9599 | 131.135 | 584.673 | X | X |
| Krauss | 44 | 0.4766 | 6.2066 | 79.102 | 287.406 | | |
| Vogler | 4 | -1.03776 | 0.9874 | 102.097 | 285.734 | X | X |
| Matsuura | 59 | -18.6596 | 1.9792 | 364.351 | 1828.691 | | |
| Taguchi_12 | 416 | 1.6921 | 39.6054 | 437.558 | 2406.648 | | |
| MTBLS36 | 31 | -7.5698 | 4.0662 | 90.078 | 504.438 | | |
| Huntscha | 18 | -1.2349 | 5.458 | 85.066 | 503.515 | X | X |
| Ales_18 | 19 | -2.86343 | 4.9434 | 152.149 | 822.953 | X | X |
| FEM_orbitrap_urine | 4 | 1.0193 | 2.6518 | 152.149 | 228.203 | X | X |
| Cao_HILIC | 28 | -5.6689 | 3.8257 | 59.068 | 427.413 | | X |
| FEM_short | 1 | 2.6932 | 2.6932 | 179.219 | 179.219 | X | X |
| 1290SQ | 4 | 1.68362 | 4.3806 | 133.194 | 306.402 | X | X |
| LIFE_new | 27 | -1.43816 | 3.9591 | 103.121 | 625.756 | | X |
| IPB_Halle | 27 | -1.3259 | 2.8083 | 146.193 | 372.37 | | X |
| MTBLS20 | 49 | -5.1426 | 3.6959 | 124.139 | 466.571 | | |
| Otto | 36 | -1.06463 | 6.1213 | 136.114 | 563.672 | | |
| Kojima | 33 | -3.3875 | 3.8441 | 203.249 | 513.504 | | |
| MTBLS4 | 24 | 3.7994 | 12.6126 | 467.584 | 836.189 | | X |
| MPI_Symmetry | 11 | -1.0565 | 4.9434 | 238.239 | 822.953 | X | X |
| Nikiforos | 35 | -0.55943 | 6.80734 | 114.173 | 916.112 | | |
| Beck | 247 | -3.0115 | 7.8493 | 74.083 | 1045.205 | | |
| MTBLS38 | 13 | -3.9585 | 5.3033 | 137.138 | 300.483 | X | X |
| UniToyama_Atlantis | 69 | -2.2026 | 6.4109 | 256.257 | 1109.307 | | |
| Matsuura_15 | 6 | -12.6413 | -6.753 | 732.686 | 1189.083 | X | X |
| Eawag_XBridgeC18 | 255 | -1.6561 | 11.919 | 124.165 | 995.189 | | |
| Stravs_22 | 92 | -0.3954 | 7.1837 | 129.159 | 785.891 | | |
| FEM_orbitrap_plasma | 3 | -0.1251 | 8.1689 | 77.152 | 516.455 | X | X |
| FEM_long | 142 | -9.7472 | 10.4322 | 59.072 | 1272.453 | | |
| PFR-TK72 | 13 | -2.6973 | 6.4126 | 154.165 | 1034.2 | X | X |
| RIKEN | 130 | -7.573 | 7.5065 | 73.095 | 932.834 | | |

*Table S-3: LC-type and used column of all data sets*

| Data set | LC-type | Column |
|---|---|---|
| Krauss | Reversed-phase | Kinetex Core-Shell C18 2.6 um, 3.0 x 100 mm, Phenomenex |
| Nikiforos | Reversed-phase | Acclaim RSLC C18 2.2um, 2.1x100mm, Thermo |
| Tohge | Reversed-phase | ACQUITY UPLC BEH C18 2.1 by 100 mm (Waters, Milford, MA,USA) |
| Otto | Reversed-phase | XBridge C18 3.5um, 2.1x50mm, Waters |
| FEM_short | Reversed-phase | Waters ACQUITY UPLC HSS T3 C18 |
| 1290SQ | Unknown | Unknown |
| LIFE_new | Reversed-phase | Waters ACQUITY UPLC HSS T3 C18 |
| LIFE_old | Reversed-phase | Waters ACQUITY UPLC BEH C18 |
| IPB_Halle | Reversed-phase | Waters ACQUITY UPLC HSS T3 C18 |
| MTBLS87 | HILIC | Merck SeQuant ZIC-pHILIC column |
| FEM_lipids | Reversed-phase | Ascentis Express C18 |
| FEM_orbitrap_plasma | Reversed-phase | Waters ACQUITY UPLC HSS T3 C18 |
| Cao_HILIC | HILIC | Merck SeQuant ZIC-pHILIC column |
| FEM_orbitrap_urine | Reversed-phase | Waters ACQUITY UPLC HSS T3 C18 |
| MPI_Symmetry | Reversed-phase | Symmetry C18 Column, Waters |
| PFR-TK72 | Reversed-phase | Symmetry C18 Column, Waters |
| MTBLS38 | Reversed-phase | Waters ACQUITY UPLC HSS T3 C18 |
| MTBLS36 | Reversed-phase | Waters ACQUITY UPLC HSS T3 C18 |
| Mark | Reversed-phase | Waters ACQUITY UPLC HSS T3 C18 |
| Toshimitsu | Reversed-phase | Waters Atlantis T3 (2.1 x 150 mm, 5 um) |
| Matsuura | HILIC | TSK-GEL Amide-80 2.0 mm X 250 mm (TOSOH) |
| UniToyama_Atlantis | Reversed-phase | Waters Atlantis T3 (2.1 x 150 mm, 5 um) |
| Ales_18 | Reversed-phase | Symmetry C18 Column, Waters |
| Matsuura_15 | Reversed-phase | Wakosil 5C18-200 2.0 mm X 250 mm (Wako) |
| Stravs | Reversed-phase | XBridge C18 3.5um, 2.1x50mm, Waters |
| Beck | Reversed-phase | XBridge C18 3.5um, 2.1x50mm, Waters |
| RIKEN | Reversed-phase | Waters ACQUITY UPLC BEH C18 |
| Takahashi | Reversed-phase | TOSOH TSKgel ODS-100V5um Part no. 21456 |
| Krauss_21 | Reversed-phase | Kinetex Core-Shell C18 2.6 um, 3.0 x 100 mm, Phenomenex |
| UFZ_Phenomenex | Reversed-phase | Kinetex Core-Shell C18 2.6 um, 3.0 x 100 mm, Phenomenex |
| FEM_long | Reversed-phase | Waters ACQUITY UPLC HSS T3 C18 |
| Taguchi | Reversed-phase | ACQUITY UPLC BEH C18 column, Waters |
| Kohlbacher | Reversed-phase | ACQUITY UPLC BEH C8 column (2.1x100 mm, 1.7 um, Waters, USA) |
| MTBLS20 | Reversed-phase | Hypersil Gold 1.9 um C18 |
| Eawag_XBridgeC18 | Reversed-phase | XBridge C18 3.5u 2.1x50 mm |
| Stravs_22 | Reversed-phase | XBridge C18 3.5um, 2.1x50mm, Waters |
| Taguchi_12 | Reversed-phase | Develosil C30, Nomura Chemical |
| Vogler | Reversed-phase | XBridge C18 3.5um 2.1x50mm Waters |
| Huntscha | Reversed-phase | XBridge C18 3.5um 2.1x50mm Waters |
| Ken | Reversed-phase | Waters Atlantis T3 (2.1 x 150 mm 5 um) |
| Kojima | Reversed-phase | ACQUITY UPLC BEH C18 2.1 by 50 mm (Waters Milford MA USA) |
| MTBLS17 | Reversed-phase | Waters ACQUITY C18 |
| MTBLS19 | Reversed-phase | Waters ACQUITY C18 |
| MTBLS39 | Reversed-phase | Alltima HP RP-C18 |
| MTBLS4 | Reversed-phase | Hypersil Gold 1.9 µm C18 |
| MTBLS52 | Reversed-phase | Waters ACQUITY UPLC BEH C18 |

*Table S-4: List of the molecular descriptors (features) calculated using RDKit that are used to train the models for $t_R$ prediction*

| | | | |
|---|---|---|---|
| fr_C_O_noCOO | PEOE_VSA12 | SlogP_VSA6 | fr_allylic_oxid |
| PEOE_VSA3 | PEOE_VSA11 | SlogP_VSA7 | VSA_EState8 |
| Chi4v | PEOE_VSA10 | SlogP_VSA1 | VSA_EState9 |
| fr_Ar_COO | BalabanJ | SlogP_VSA2 | fr_piperdine |
| fr_SH | fr_lactone | SlogP_VSA3 | fr_Ar_OH |
| Chi4n | fr_Al_COO | NumRadicalElectrons | fr_sulfide |
| SMR_VSA10 | EState_VSA11 | fr_NH2 | NOCount |
| fr_para_hydroxylation | Chi0 | fr_piperzine | Chi1n |
| fr_Ar_NH | Chi1 | NumHeteroatoms | PEOE_VSA8 |
| fr_halogen | NumAliphaticRings | fr_NH1 | PEOE_VSA7 |
| fr_dihydropyridine | MolLogP | MaxAbsEStateIndex | PEOE_VSA6 |
| fr_priamide | fr_nitro | fr_amide | PEOE_VSA4 |
| SlogP_VSA4 | NumAliphaticCarbocycles | Chi3n | MaxEStateIndex |
| fr_guanido | fr_C_O | SMR_VSA3 | PEOE_VSA2 |
| MinPartialCharge | fr_ether | SMR_VSA1 | PEOE_VSA1 |
| fr_furan | fr_alkyl_halide | Chi3v | NumSaturatedCarbocycles |
| NumAromaticCarbocycles | NumValenceElectrons | SMR_VSA6 | Chi1v |
| fr_COO2 | fr_aryl_methyl | Kappa3 | fr_Al_OH_noTert |
| fr_amidine | fr_Ndealkylation2 | Kappa2 | fr_epoxide |
| SMR_VSA7 | MinEStateIndex | EState_VSA6 | fr_hdrzone |
| fr_benzodiazepine | fr_term_acetylene | EState_VSA7 | fr_isothiocyan |
| fr_Imine | HallKierAlpha | SMR_VSA9 | fr_phenol |
| MolWt | fr_C_S | EState_VSA5 | MolMR |
| fr_hdrzine | fr_ketone_Topliss | EState_VSA2 | PEOE_VSA9 |
| fr_urea | Ipc | EState_VSA3 | fr_aldehyde |
| NumAromaticRings | fr_HOCCN | fr_Ndealkylation1 | fr_pyridine |
| fr_quatN | fr_phos_ester | EState_VSA1 | fr_tetrazole |
| NumAliphaticHeterocycles | BertzCT | fr_ketone | fr_nitro_arom_nonortho |
| fr_benzene | SlogP_VSA12 | SMR_VSA5 | Chi0v |
| fr_phos_acid | EState_VSA9 | MinAbsEStateIndex | NumRotatableBonds |
| fr_sulfone | SlogP_VSA11 | fr_Ar_N | MaxAbsPartialCharge |
| VSA_EState10 | fr_COO | fr_Nhpyrrole | |
| fr_aniline | NHOHCount | fr_ester | |
| fr_N_O | fr_unbrch_alkane | EState_VSA4 | |
| fr_sulfonamd | NumSaturatedRings | NumHDonors | |
| fr_thiazole | MaxPartialCharge | fr_oxime | |
| TPSA | fr_methoxy | Chi2v | |
| EState_VSA8 | fr_thiophene | HeavyAtomCount | |
| PEOE_VSA14 | MinAbsPartialCharge | NumHAcceptors | |
| PEOE_VSA13 | SlogP_VSA5 | fr_lactam | |

*Table S-5: List of molecular descriptors (features) calculated using RDKit that were excluded for model training based on high correlation with other features ($R^2 > 0.96$) or low standard deviation ($stdev <= 0.01$).*

| | | | |
|---|---|---|---|
| fr_barbitur | VSA_EState7 | fr_diazo | PEOE_VSA5 |
| ExactMolWt | VSA_EState1 | SMR_VSA4 | fr_imide |
| NumSaturatedHeterocycles | VSA_EState2 | VSA_EState5 | FractionCSP3 |
| EState_VSA10 | VSA_EState3 | fr_prisulfonamd | NumAromaticHeterocycles |
| HeavyAtomMolWt | SlogP_VSA10 | SMR_VSA8 | fr_bicyclic |
| fr_nitro_arom | SlogP_VSA8 | fr_isocyan | Kappa1 |
| fr_phenol_noOrthoHbond | SlogP_VSA9 | Chi2n | Chi0n |
| fr_thiocyan | fr_NH0 | fr_azide | RingCount |
| VSA_EState4 | LabuteASA | fr_oxazole | fr_ArN |
| VSA_EState6 | SMR_VSA2 | fr_alkyl_carbamate | |
| fr_Al_OH | fr_azo | fr_nitrile | |
| PEOE_VSA5 | fr_morpholine | fr_nitroso | |

*Listing S-1: Distributions of hyperparameters used to optimize the regression models*

```
# LASSO

params = {
"alpha" : uniform(0.0,10.0),
"copy_X" : [True],
"fit_intercept" : [True],
"normalize" : [False],
"precompute" : [True,False],
"max_iter" : [5000]
}

# AB

params = {
"n_estimators": randint(10,100), #,1000,3000], #
"learning_rate": uniform(0.01,0.5)
}

# GB

params = {
"n_estimators" : randint(20,100),
"max_depth" : randint(1,12),
"learning_rate" : uniform(0.01,0.25),
"gamma" : uniform(0.0,10.0),
"reg_alpha" : uniform(0.0,10.0),
"reg_lambda" : uniform(0.0,10.0)
}

# SVR

params = {
"epsilon": uniform(0.01,100.0),
"kernel": ["linear","rbf"],
"degree": randint(1,12),
"gamma" : uniform(0.000001,100),
}

# BRR

params = {
"n_iter" : [50],
"alpha_1" : uniform(1e-10,1e-2),
"lambda_1" : uniform(1e-10,1e-2),
"threshold_lambda" : randint(1,10000),
"verbose" : [True]
}
```

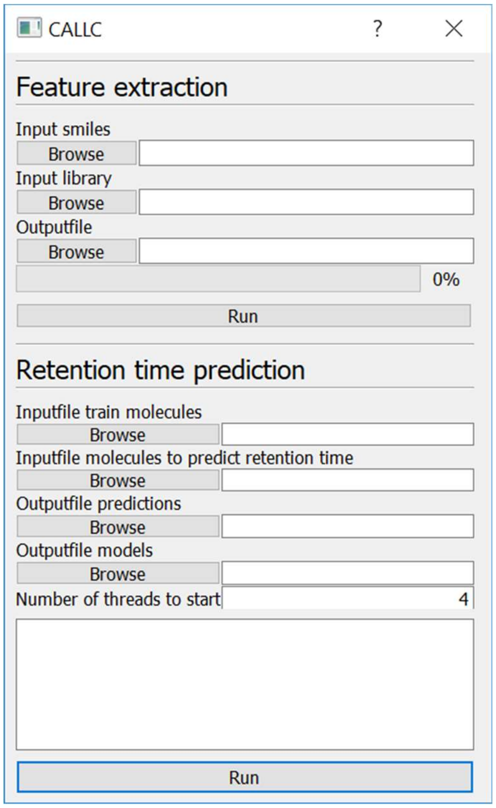

*Figure S-1: The Graphical User Interface for CALLC to predict the retention time for new data sets. A user manual and code are available at: https://github.com/RobbinBouwmeester/CALLC*

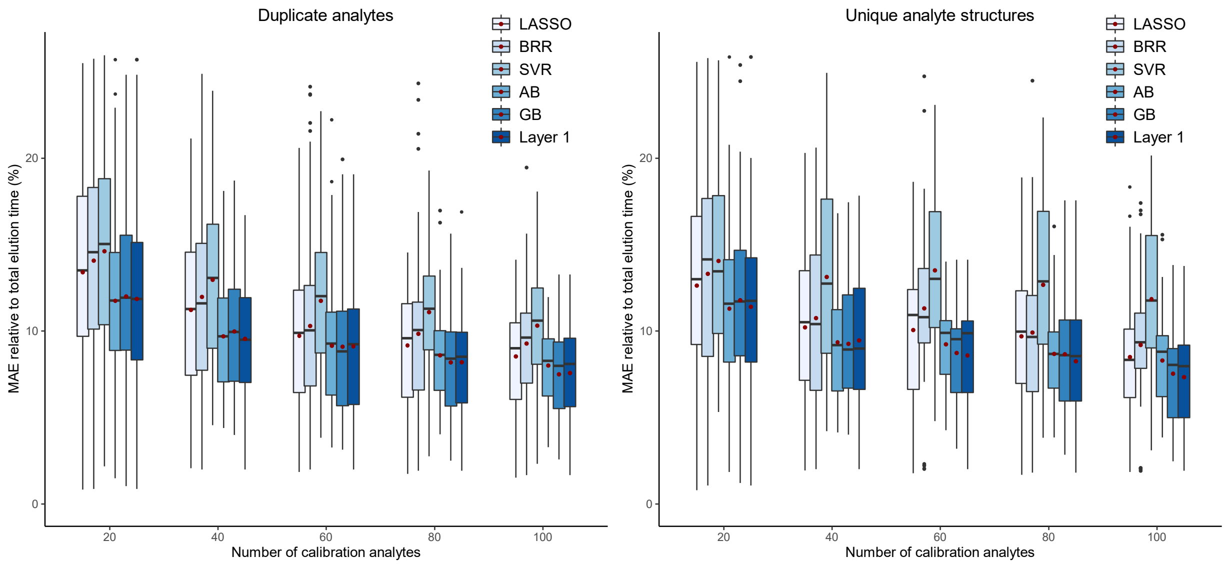

*Figure S-2: Comparison of different regression models in Layer 1 and the model predictions that were selected based on the CV performance (labeled as Layer 1). The evaluation metric is the Relative Mean Absolute Error (RMAE) and the red dot shows the mean value of this metric. The left panel consists of 34 data sets that have shared analyte structures between data sets. The right panel consists of 21 data sets that do not share any analyte structures between data sets.*

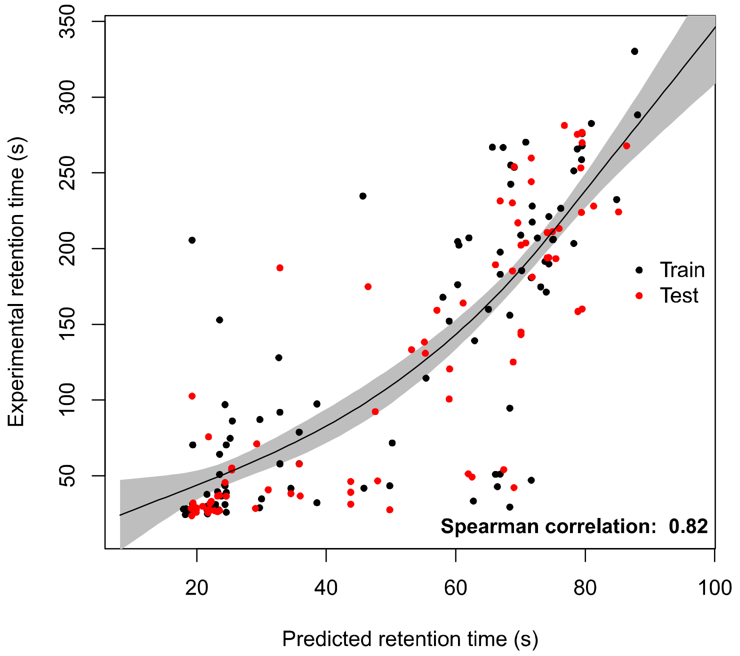

*Figure S-3: Example of a GAM model that is used to calibrate predictions (in Layer 2) from a model based on the data set RIKEN to the data set LIFE_new. The black points show the predictions used for fitting the calibration curve and the red points that are part of the test set. The shaded grey area is the standard deviation of the fit.*

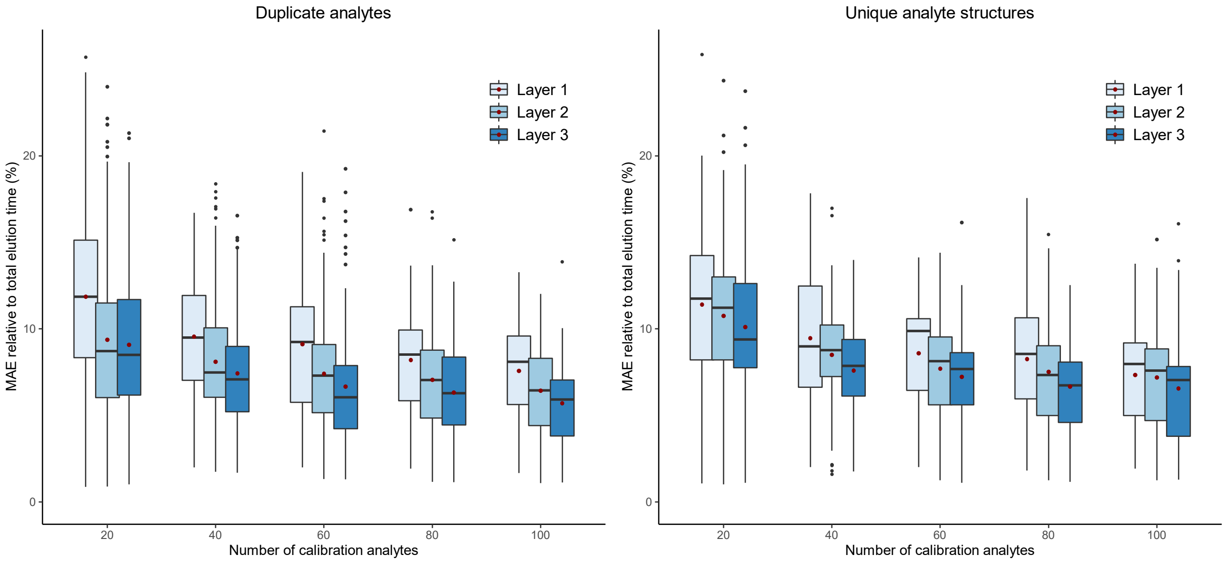

*Figure S-4: Performance evaluation of Layer 1, Layer 2, and Layer 3. The evaluation metric is the Relative Mean Absolute Error (RMAE) and the red dot shows the mean value of this metric. The left panel consists of 34 data sets that have shared analyte structures between data sets. The right panel consists of 21 data sets that do not share any analyte structures between data sets.*

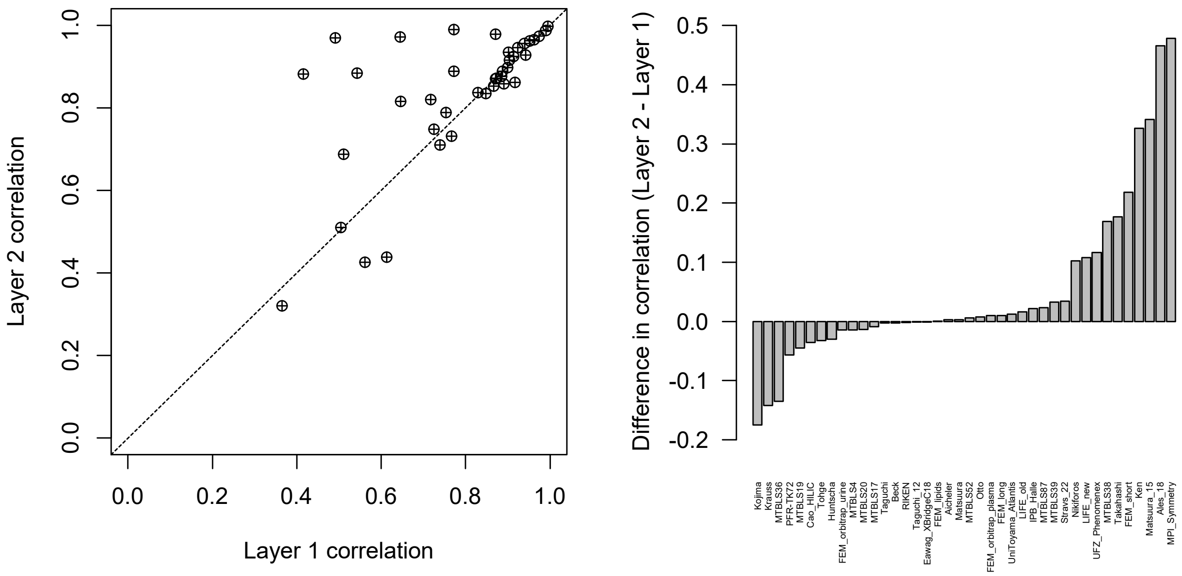

*Figure S-5: CV performance evaluation between Layer 1 and Layer 2 on 40 data sets, where shared analytes structures between data sets are allowed. The evaluation metric is the Pearson correlation between predicted and observed retention times. The left panel shows the achieved correlation for each data set in both layers, where the dotted line indicates the position where both layers perform equally. The right panel shows the difference in the Pearson correlation between the layers. Positive values mean that Layer 1 had a higher correlation than Layer 2, with the height of the bar showing the magnitude of the difference between the correlation values. Negative values show a higher correlation in Layer 1 than Layer 2. One data point, for Krauss, in the left panel is outside of the plotting area.*

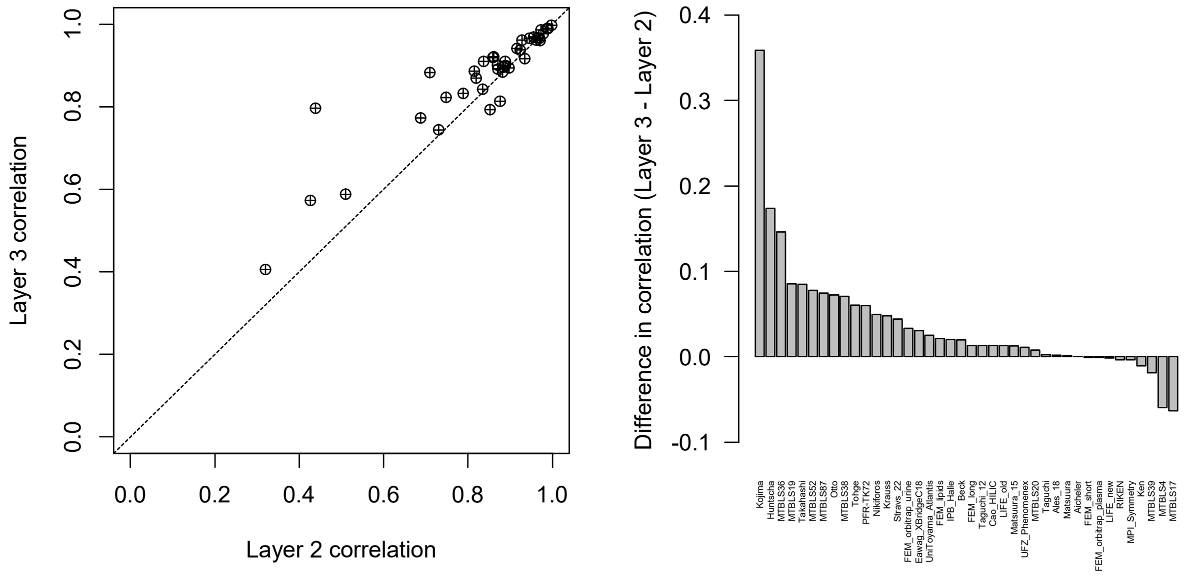

*Figure S-6: CV performance evaluation between Layer 2 and Layer 3 on 40 data sets, where shared analytes structures between data sets are allowed. The evaluation metric is the Pearson correlation between predicted and observed retention times. The left panel shows the achieved correlation for each data set in both layers, where the dotted line indicates the position where both layers perform equally. The right panel shows the difference in the Pearson correlation between the layers. Positive values mean that Layer 3 had a higher correlation than Layer 2, with the height of the bar showing the magnitude of the difference between the correlation values. Negative values show a higher correlation in Layer 2 than Layer 3. One data point, for Krauss, in the left panel is outside of the plotting area.*

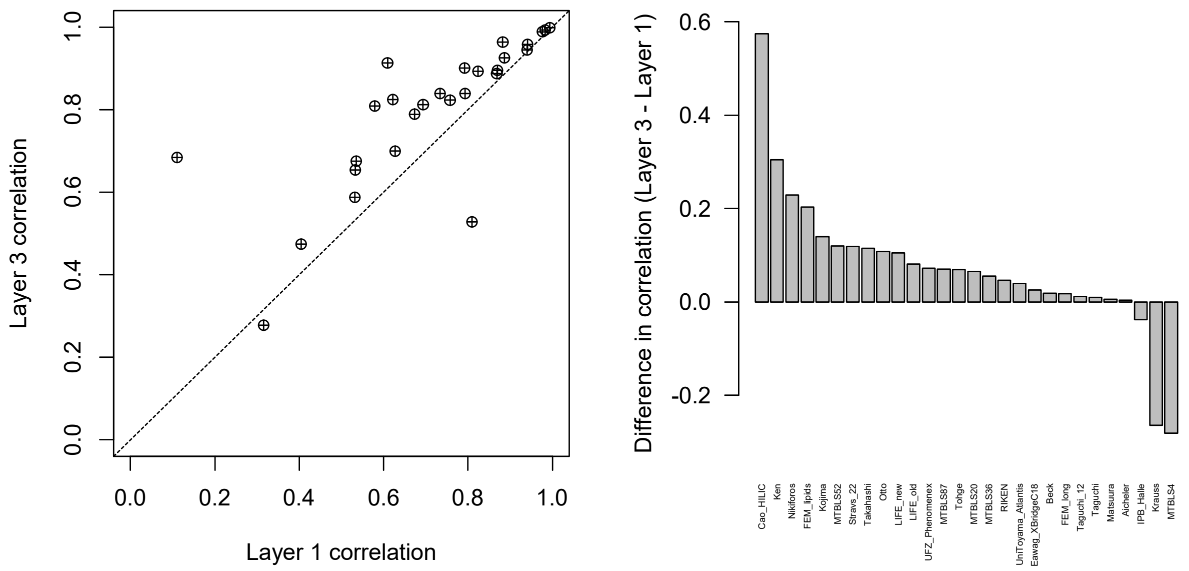

*Figure S-7: CV performance evaluation between Layer 1 and Layer 3 on 28 data sets, where shared analytes structures between data sets are removed. The evaluation metric is the Pearson correlation between predicted and observed retention times. The left panel shows the achieved correlation for each data set in both layers, where the dotted line indicates the position where both layers perform equally. The right panel shows the difference in the Pearson correlation between the layers. Positive values mean that Layer 3 had a higher correlation than Layer 1, with the height of the bar showing the magnitude of the difference between the correlation values. Negative values show a higher correlation in Layer 1 than Layer 3. One data point, for Krauss, in the left panel is outside of the plotting area.*

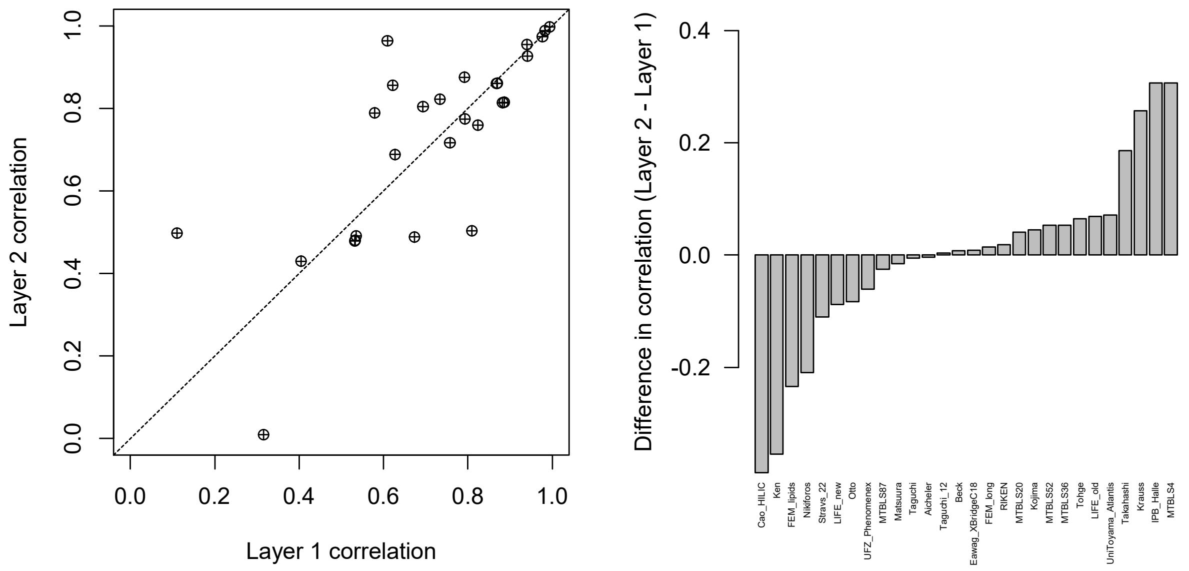

*Figure S-8: CV performance evaluation between Layer 1 and Layer 2 on 28 data sets, where shared analytes structures between data sets are removed. The evaluation metric is the Pearson correlation between predicted and observed retention times. The left panel shows the achieved correlation for each data set in both layers, where the dotted line indicates the position where both layers perform equally. The right panel shows the difference in the Pearson correlation between the layers. Positive values mean that Layer 2 had a higher correlation than Layer 1, with the height of the bar showing the magnitude of the difference between the correlation values. Negative values show a higher correlation in Layer 1 than Layer 2.*

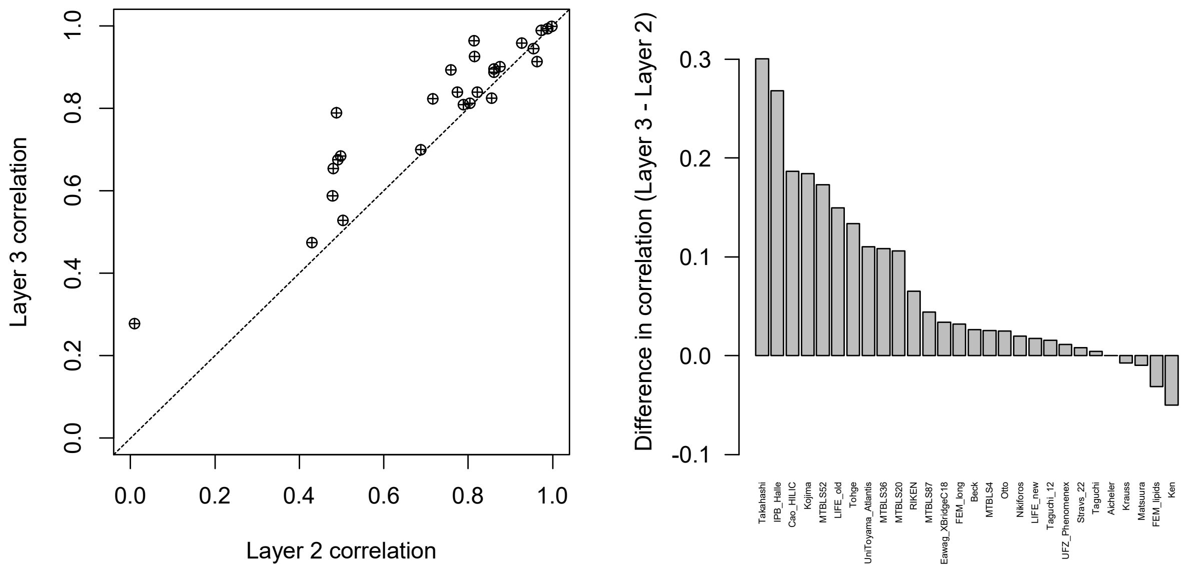

*Figure S-9: CV performance evaluation between Layer 2 and Layer 3 on 28 data sets, where shared analytes structures between data sets are removed. The evaluation metric is the Pearson correlation between predicted and observed retention times. The left panel shows the achieved correlation for each data set in both layers, where the dotted line indicates the position where both layers perform equally. The right panel shows the difference in the Pearson correlation between the layers. Positive values mean that Layer 3 had a higher correlation than Layer 2, with the height of the bar showing the magnitude of the difference between the correlation values. Negative values show a higher correlation in Layer 2 than Layer 3. One data point, for Krauss, in the left panel is outside of the plotting area.*

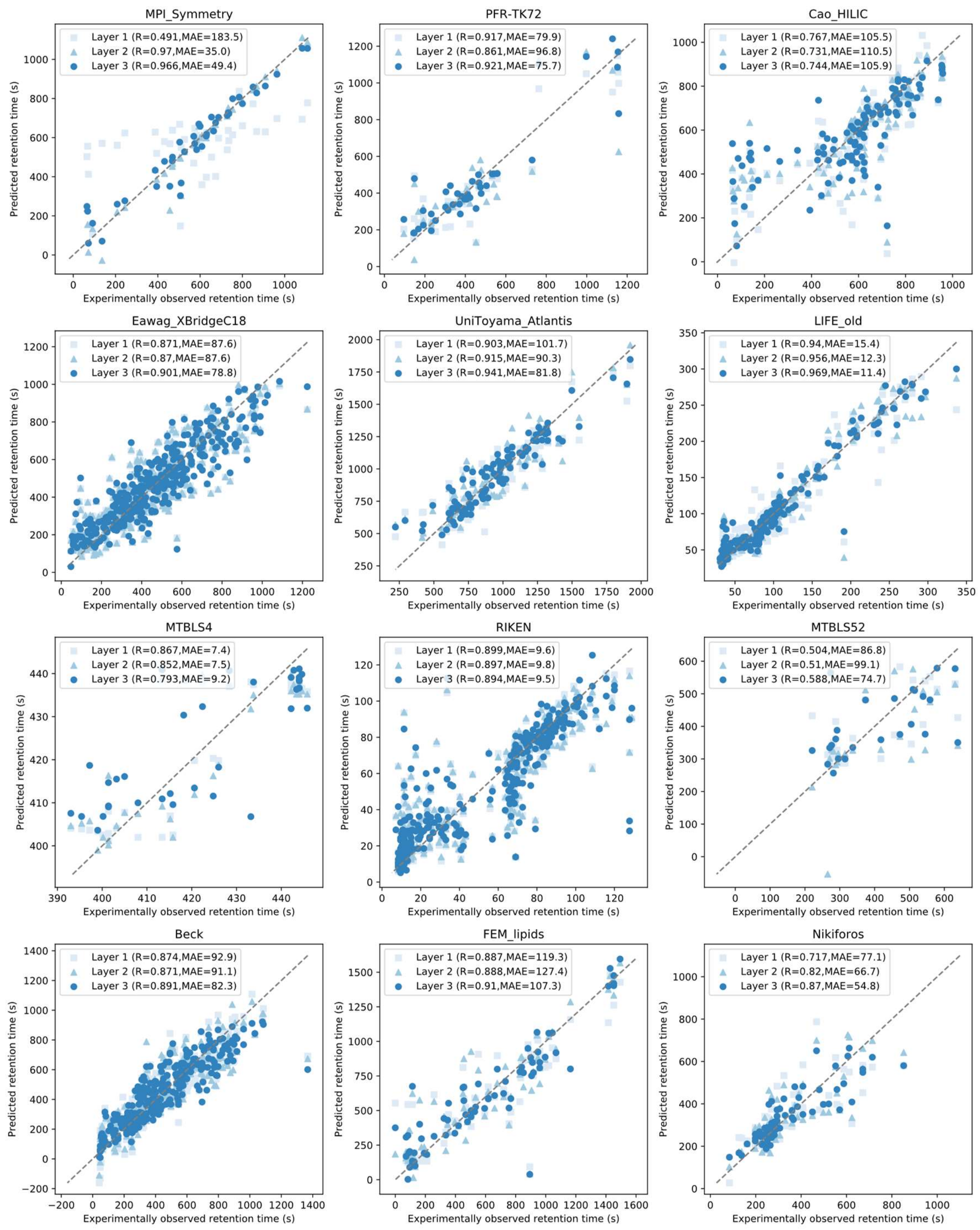

*Figure S-10: Scatter plots between the predicted and observed retention times for 9 data sets where shared analytes structures between data sets are allowed. The different shades of blue show the predictions made by the different layers in CALLC.*

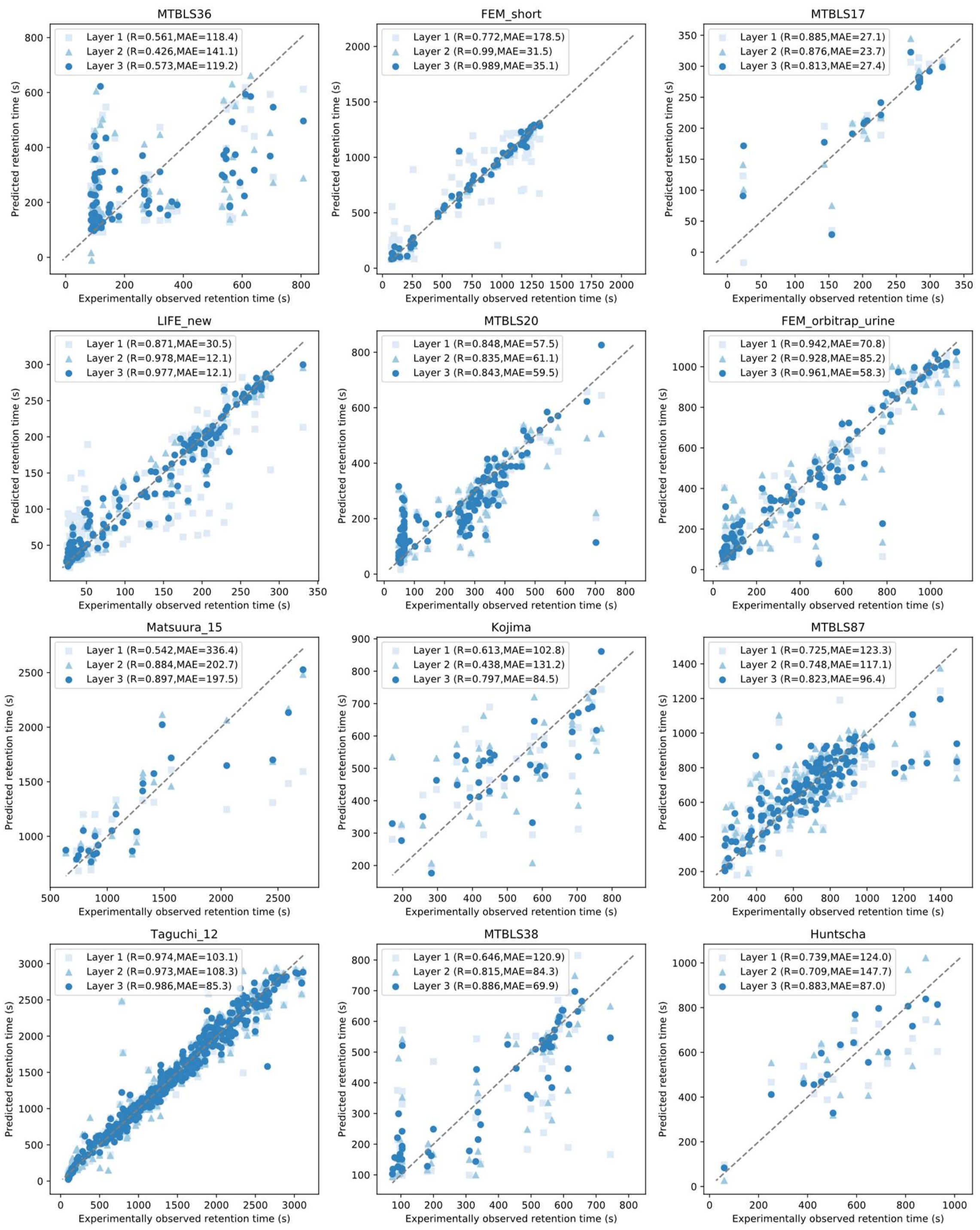

*Figure S-11: Scatter plots between the predicted and observed retention times for 9 data sets where shared analytes structures between data sets are allowed. The different shades of blue show the predictions made by the different layers in CALLC.*

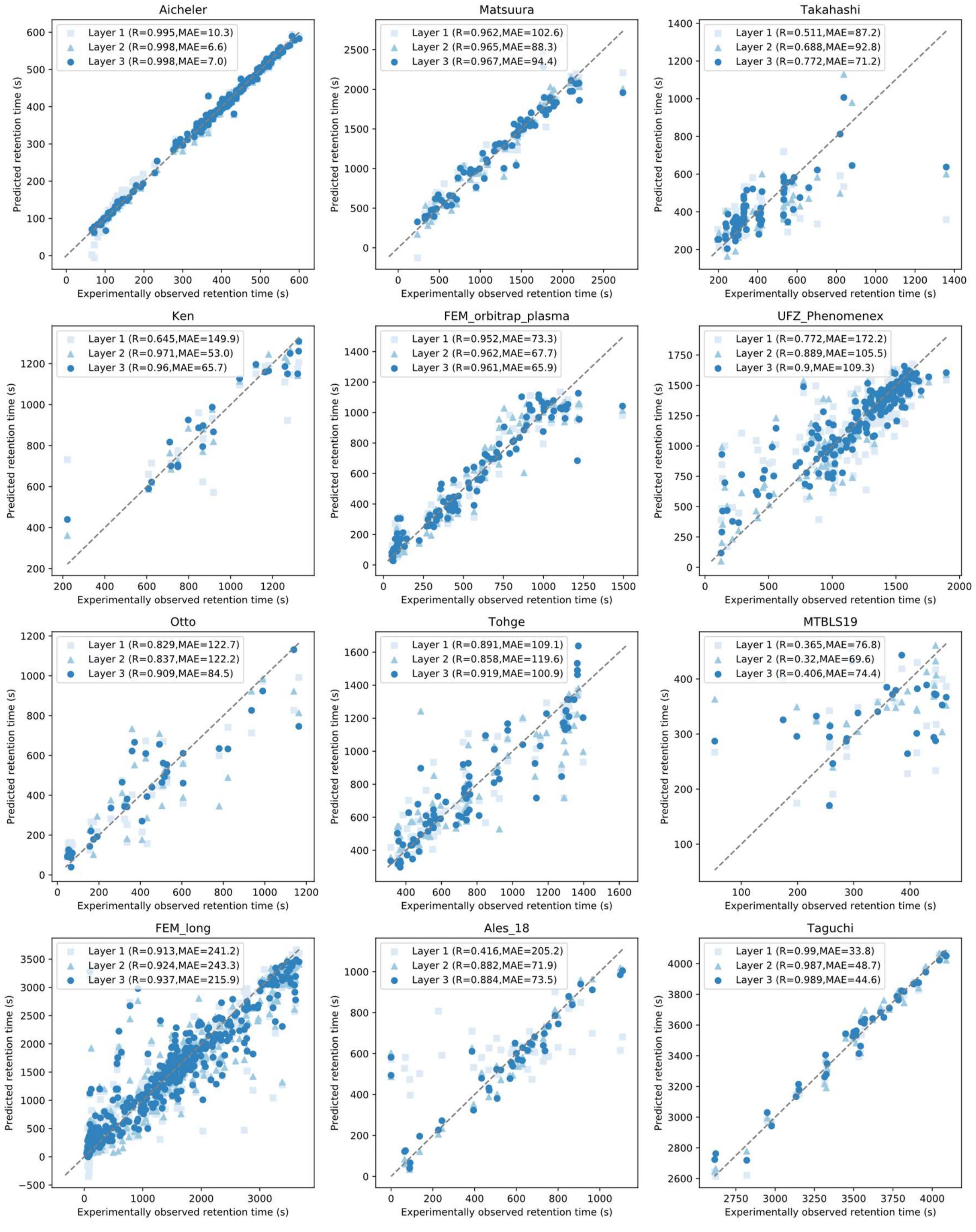

*Figure S-12: Scatter plots between the predicted and observed retention times for 9 data sets where shared analytes structures between data sets are allowed. The different shades of blue show the predictions made by the different layers in CALLC.*

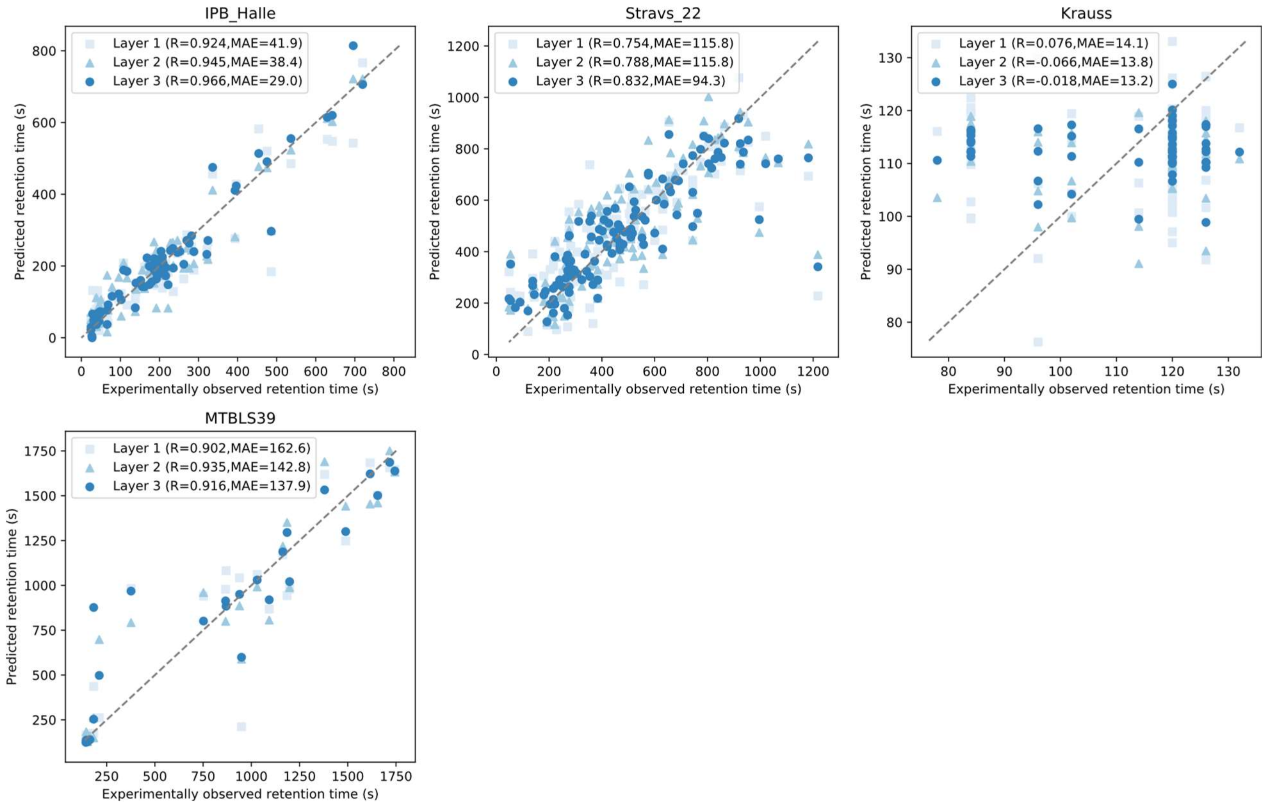

*Figure S-13: Scatter plots between the predicted and observed retention times for 4 data sets where shared analytes structures between data sets are allowed. The different shades of blue show the predictions made by the different layers in CALLC.*

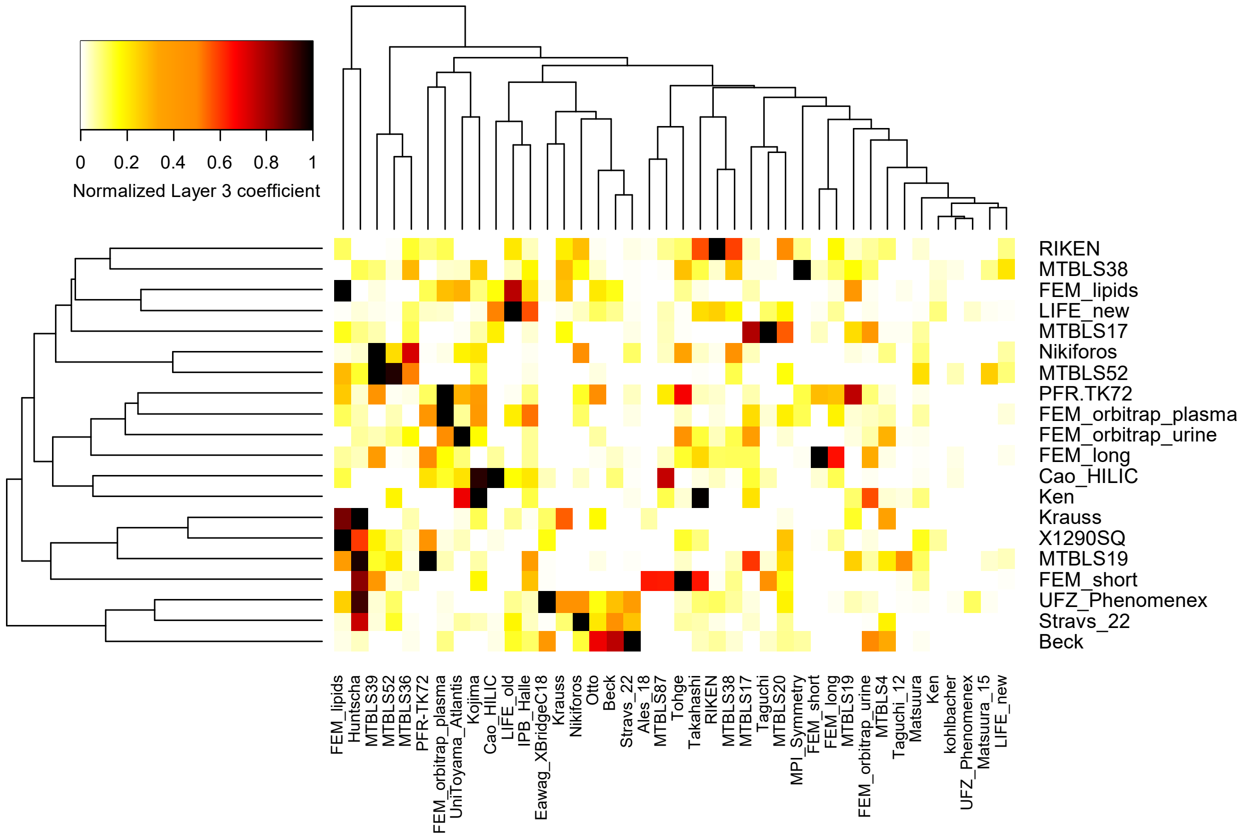

*Figure S-14: The normalized layer 3 coefficients for different data sets (rows) and the utilized regression models (columns). The coefficients are summed among all initial training sizes and different regression models. The summed coefficients are divided by the max coefficient. Only those prediction sets that have a summed coefficient in the top 20 are displayed.*

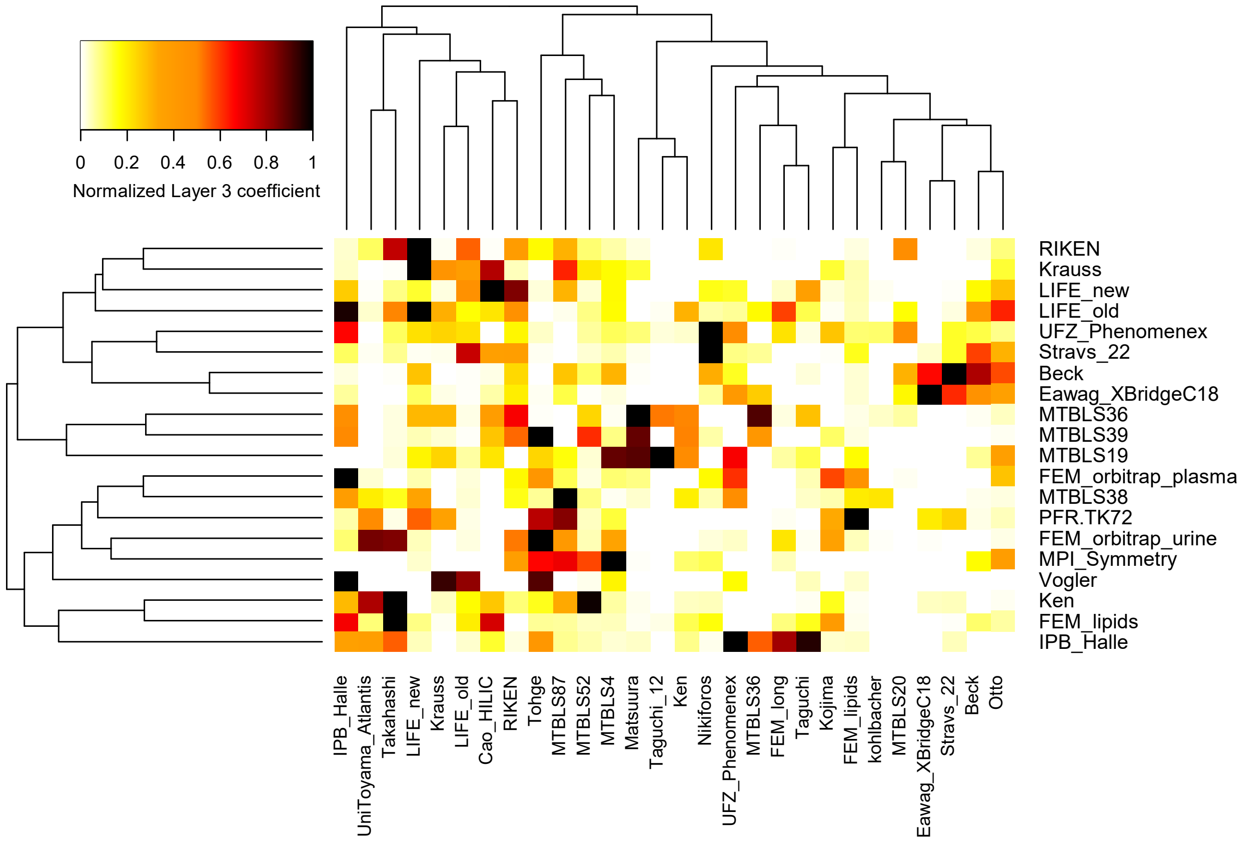

*Figure S-15: The normalized layer 3 coefficients for different data sets (rows) and the utilized regression models (columns). The coefficients are summed among all initial training sizes and different regression models. The summed coefficients are divided by the max coefficient. Only those prediction sets that have a summed coefficient in the top 20 are displayed.*
